# ASpli2: Integrative analysis of splicing landscapes through RNA-Seq assays

**DOI:** 10.1101/2020.06.21.162891

**Authors:** Estefania Mancini, Andres Rabinovich, Javier Iserte, Marcelo Yanovsky, Ariel Chernomoretz

**Author notes:** Equally contributed (Estefania Mancini, Andres Rabinovich).

## Abstract

Genome-wide analysis of alternative splicing has been a very active field of research since the early days of NGS (Next generation sequencing) technologies. Since then, ever-growing data availability and the development of increasingly sophisticated analysis methods have uncovered the complexity of the general splicing repertoire. However, independently of the considered quantification methodology, very often changes in variant concentration profiles can be hard to disentangle. In order to tackle this problem we present ASpli2, a computational suite implemented in R, that allows the identification of changes in both, annotated and novel alternative splicing events, and can deal with complex experimental designs.

Our analysis workflow relies on the analysis of differential usage of subgenic features in combination with a junction-based description of local splicing changes. Analyzing simulated and real data we found that the consolidation of these signals resulted in a robust proxy of the occurrence of splicing alterations. While junction-based signals allowed us to uncover annotated as well and non-annotated events, bin-associated signals notably increased recall capabilities at a very competitive performance in terms of precision.

## 1. Introduction

The vast majority of protein coding genes in eukaryotic organisms are transcribed into precursor RNA messenger molecules (pre-mRNA) carrying protein coding regions (exons) interleaved by non-coding ones (introns). The later are removed in a co-transcriptional dynamical maturation process called splicing. Alternative splicing (AS) occurs whenever distinct splicing sites are selected in this process resulting in different mature mRNA molecules [1, 2].

Far from being an exception, it was found that AS is a rather common mechanism of gene regulation that serves to expand the functional diversity of a single gene allowing the generation of multiple mRNA isoforms from a single genomic locus [3]. Five basic modes of AS are generally recognized: the skipping of a given exon (exon skipping, ES), the exon elongation/contraction produced by the use of alternative 5’ donor (Alt5’) or 3’ acceptor (Alt3’) sites respectively, the retention of an intronic stretch in the mature mRNA form (intron retention IR), and the alternative use of mutually exclusive exons (MEX). These canonical forms of AS are prevalent among eukaryotes, although their relative incidence might vary between them [4]. Despite their ubiquity, these simple patterns that mainly involve binary choices of exons, donor and acceptor sites, do not exhaust the splicing repertoire. On the contrary, much more complex biologically relevant patterns could arise [5, 6]. In practice the study of AS faces many technical challenges that cause that every quantitative approach typically suffers methodological limitations. Despite the use of different statistical approaches, some methods consider only preexisting known annotation, some can exclusively handle canonical splicing events and some can only handle pairwise comparisons between conditions (for a comprehensive review see [7, 8, 9]).

The analysis of AS at genomic scale started-in with microarray technologies [10, 11] and nowadays is routinely probed using RNAseq assays [12, 13]. Roughly speaking, there are three main computational approaches to study splicing diversity from RNAseq data. For one hand there are transcript reconstruction methods, like MISO [14] or Cufflink [15] that aim to infer a probabilistic model of the frequency of each isoform from the read distribution mapped to a given gene. In the same spirit, Kallisto [16] and Salmon[17] are two recently introduced methods that leverage on light-weight pseudoalignment heuristics to quantify transcript abundances. For the other hand, methods like DEXSeq [18], edgeR [19, 20], or voom-limma [21], focus on the analysis of differential usage of subgenic features (e.g. exons) between conditions. Finally, there are also methods like rMATS [22], MAJIQ [5] or LeafCutter [23] that leverage on junction information to infer both, annotated and novel splicing events.

Differently form other approaches, ASpli2 was specifically designed to integrate several independent signals in order to deal with the complexity that might arise in splicing patterns. Taking into account genome annotation information, ASpli2 considers bin-based signals along with junction inclusion indexes in order to assess for statistically significant changes in read coverage. In addition, annotation-independent signals are estimated based on the complete set of experimentally detected splice junctions. Noticeably, ASpli2 can provide a comprehensive description of genome-wide splicing alterations even for non-trivial experimental designs. Our approach relies on a generalized linear model framework (as implemented in edgeR R-package [24]) to assess for the statistical analysis of specific contrasts of interest.

In order to weigh ASpli2’s performance we compared it against three different state-of-the-art methodologies: rMATS [22], LeafCutter [23] and MAJIQ [5]. The first one is a widely used piece of software that can integrate coverage and junction information to assess for changes in splicing patterns. Additionaly, LeafCutter and MAJIQ are two recently introduced methodologies that are widely used by the bioinformatics community. Both approaches focus on the analysis of clusters of junctions to study local splicing patterns of varying complexity. However, they differ in many technical and statistical aspects [5]. For instance, LeafCutter was not designed to handle intron retention events and considers a Dirichlet-multinomial generalized linear model to test for differential intron excision between two groups of samples. MAJIQ, on the other hand, relies on a bayesian estimation of the posterior Percent Selected Index to identify splicing affected junctions.

Other methodolgies like DEXSeq [18], edgeR [24], or voom [21] are also widely considered for bioinformatics analysis as they are very versatile tools to analyse differential usage of exons from RNA-seq data. In fact, ASpli2 makes use of the genome-binning scheme originally presented in DEXseq to quantify read coverage signals (see Sup.Mat.8.1) and leverages on the statistical framework developed in edgeR to estimate robust splicing signals (see Section 3.1). These methodologies constitute great toolboxes to implement ad-hoc analysis. However, as they do not intend to provide *per se* selfcontained solutions that produces final reports starting from read-alignment input data they were not explicitly included in our analysis.

The paper is organized as follows. In Section 2.2 we analyzed a simulated dataset in order to evaluate the specificity and sensitivity of ASpli2 discoveries. These results were contrasted against LeafCutter, MAJIQ and rMATS outcomes in order to analyse ASpli2 performance. In Section 2.3 we explored the ability of ASpli2 to uncover consistent splicing-patterns from two independent RNAseq assays that probed the same biology. We focused on the alterations of splicing patterns of *A. thaliana* transcriptome caused by the knock out of PRMT5, a methyl transferase that, among other proteins, targets several Sm spliceosomal proteins [25, 26, 27]. This analysis was also performed with the other considered methodologies in order to compare their capacity to generate reproducible results. In addition, we capitalized on ASpli2’s ability to handle complex experimental designs to produce a consolidated data-set from the independent assays. In this section we also aimed to quantify the level of agreement of ASpli2, LeafCutter, MAJIQ and rMATS discoveries with qRT-PCR based alternative splicing evidence. To that end, we took advantage of two independent studies that analyzed splicing altered events in PRMT5 mutants using qRT-PCR assays [25, 28]. Finally, in Section 2.4, we considered a 28 samples paired-study of human prostate cancer [29]. Using this data-set we analyzed how the performance, time and memory requirements scaled with the number of considered samples in a paired experimental design. Finally, we discussed our results in Section 4 and presented our conclusions in Section 5.

## 2. Results

### 2.1. ASpli2 workflow

ASpli2 was designed as a flexible R package to carry out all the major tasks required for gene expression and splicing analysis. A typical ASpli2 workflow involves: parsing the genome annotation into subgenic features called bins, overlapping read alignments against them, perform junction counting, fulfill inference tasks of differential bin and junction usage and, finally, report integrated splicing signals. A workflow diagram and a summary of ASpli2 core functionality can be found as Supplementary Figures S1 and S2 respectively. As shown in Figure S1, at every step ASpli2 generates self-contained outcomes that, if required, can be easily exported and integrated into other processing pipelines. Supplementary Figure S3 shows an example of the interactive html report generated as a final output. A detailed description of ASpli2 functionality is included in ASpli2’s R vignette, which is provided as Supplementary Material.

### 2.2. Synthetic dataset

Changes in splicing patterns were simulated in a treatment vs control setup for genes of the chromosome-one of the *Arabidopsis thaliana* plant genome (three samples per condition). In our simulations, the differential usage of splicing variants affected 2451 genomic bins in 915 genes (see Material and Methods 3.3).

The ASpli2 analysis pipeline provided three different cues to probe splicing occurrence. Statistically significative evidence is collected from: bin coverage differential signals, junction anchorage changes and variations inside junction clusters (see Material and Methods 3.1). We considered bin-coverage signals with statistically significant differential coverage changes (fdr*<* 0.05) that presented either a larger than three-fold coverage fold-change or, alternatively, a change in bin-supporting junction inclusion indices larger than 0.2. For junction based signals, on the other hand, locale and anchorage indices were required to present statistically significant changes (fdr*<* 0.01) and also should display usage signal variations larger than a 0.3 level (see Material and Methods 3.2)

In Table 1 we reported the number of correctly detected simulated events, number of false positives and number of events exclusively detected by each kind of signal: bin-coverage, junction-locale and junction-anchorage. Overlaps between discoveries reported by each kind of signal were graphically reported in panel (A) of Figure 1.

**Table (1).** Splicing detection performance of the three different ASpli2 signals. True positive and false positive calls are shown in the second and third columns respectively. The number of specific discoveries exclusively reported by each signal is reported between brackets.

| ASpli2 signal | TP | FP |
| --- | --- | --- |
| bin coverage | 924 (399) | 42 |
| junction locale | 393 (35) | 2 |
| junction anchorage | 182 (15) | 6 |
| overall | 974 | 48 |

**Figure (1).**
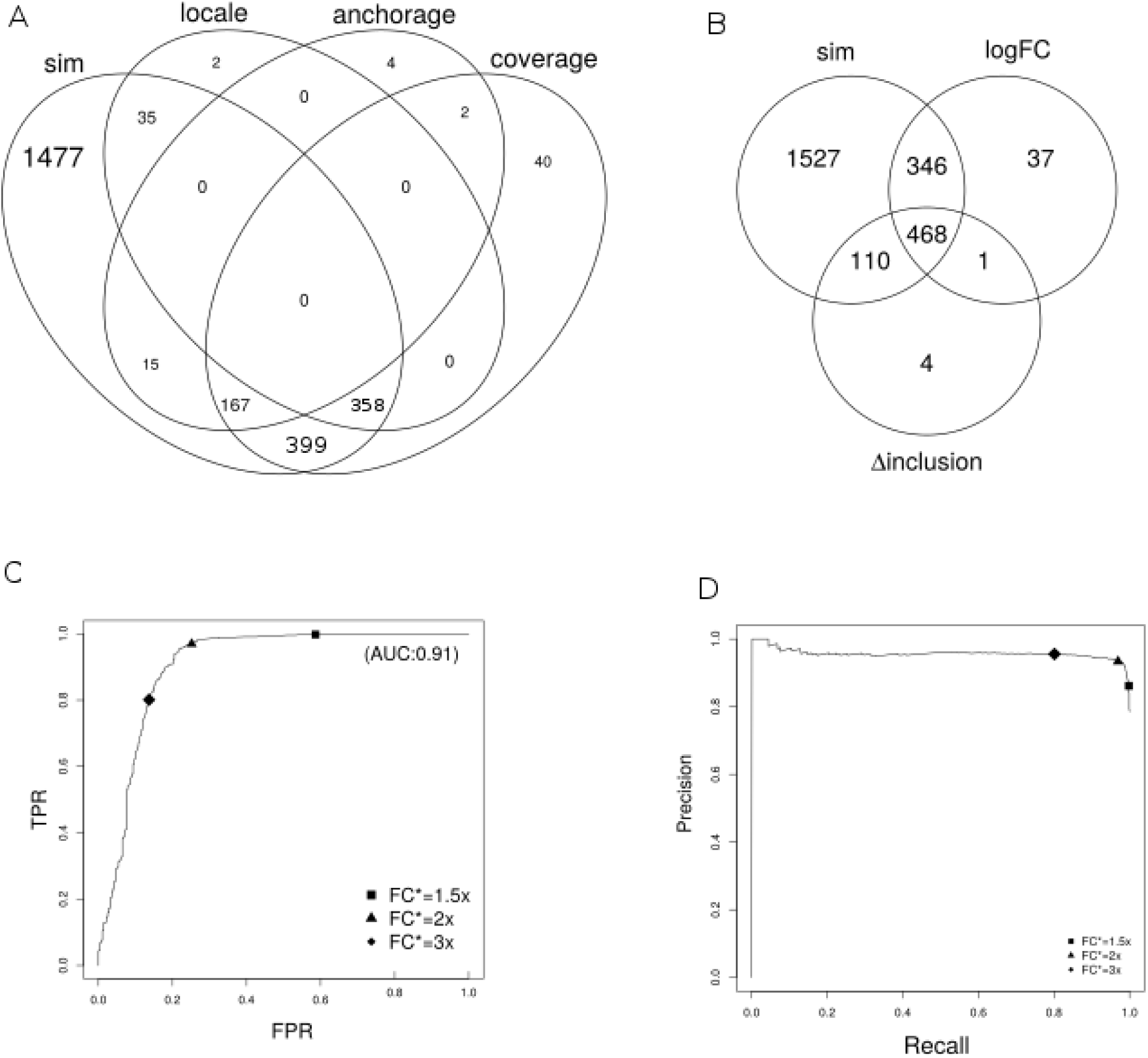
(a) Distribution of detection call produced by different ASpli2 signals. (b) Graphical summary of bin-coverage detection calls. The *sim* set correspond to simulated events. *logFC* and *D-inclusion* sets are associated to statistically significant discoveries presenting large enough fold change and large bin-supporting junction inclusion signals respectively. ROC and Precision-Recall curves, parameterized by the considered fold-change threshold level, are shown for statistically significant bins in panels (c) and (d) respectively.

It can be seen that ASpli2 correctly uncovered 974 (40%) of the 2451 simulated bin events. Moreover, we found that most of the ASpli2 undetected simulated events (1341 out of 1477) took place in genes that did not present enough expression levels over the analysed conditions and therefore were filtered out before any statistical testing (see 3.2). In fact, only 136 out of the 1110 events (12%) that did pass the gene-expression pre-filtering step were found to be false negative cases.

About 95% of ASpli2 true discoveries were identified by the analysis of significant changes at the bin-coverage level. Junction-based detection, on the other hand, could correctly identified 574 simulated events (60% of true discoveries). The null overlap between locale and anchorage detection illustrated that they probed complementary aspects of splicing events. Additionally, it can be appreciated that 41% (399) of the true discoveries were only detected by bin-coverage signals, whereas junction-based analysis contributed only 5% (50) of specific detections. A graphical summary of the detection signal landscape can be appreciated in panel (A) of Figure 1.

We decided to further characterized some aspects of bin-coverage detection calls, as this signal provided the major number of discoveries. It can be seen in panel-(B) of Figure 1 that fold-change and junction-support signals used in the bin-coverage analysis reported relevant and non-redundant information. Whereas the first one accounted for 37% of true positive instances exclusively detected by this signal, the second one accounted for the specific identification of 12% of the total number of true events. The impact of the selected fold-change threshold value, FC*, on specificity, precision and recall can be appreciated with the aid of the Receiver-Operator and Precision-Recall curves shown in panels (C) and (D) of Figure 1. It can be recognized from these figures that with the adopted 3-fold threshold ASpli2 achieved high recall and precision levels (∼ 80% and ∼ 95% respectively) laying at rather moderate levels of false positive rates (∼ 14%).

In Table 2 we reported the detection performance of ASpli2 and the results obtained by other state-of-the-art algorithms (see Sup Mat 8.2 and 8.3 for calculation details). Precision and recall values estimated at gene-level (in which a gene was reported as a discovery whenever at least one alternativesplicing event was detected within its genomic range) were reported between parenthesis. ASpli2 outcomes considering only coverage signal or just junction signals were included in the table as ASpli2_*c*_ and ASpli2_*j*_ rows respectively.

**Table (2).** Number of discoveries, precision and recall levels are reported for different detection methodologies. *ASpli*_*c*_ and *ASpli*_*j*_ correspond to ASpli2 discoveries detected using just coverage or just junction signals respectively. Values between parenthesis report quantities estimated at gene-level.

| method | size | precision | recall |
| --- | --- | --- | --- |
| ASpli2 | 1022 (631) | 0.95 (0.99) | 0.40 (0.68) |
| ASpli2<sub>c</sub> | 966 (591) | 0.96 (0.99) | 0.38 (0.64) |
| ASpli2<sub>j</sub> | 583 (456) | 0.99 (0.99) | 0.23 (0.50) |
| LeafCutter | 204 (163) | 0.93 (0.91) | 0.08 (0.16) |
| MAJIQ | 538 (381) | 0.84 (0.87) | 0.18 (0.36) |
| rMATS | 405 (352) | 0.87 (0.91) | 0.14 (0.35) |

It can be seen from the table that even though all tested algorithm shown rather high precision values, ASpli2 benefited from larger recall scores than any other methodology. Moreover, it can be appreciated that ASpli2_*j*_ displayed only marginally larger recall levels than other methodologies implying that ASpli2 leveraged on coverage signals to increase this figure of merit. All of these results suggested that ASpli2 was capable of reliably detect the simulated splicing events achieving notably high recall values at very competitive levels of precision and specificity.

### 2.3. Reproducibility Analysis

As we mentioned in the introduction, PRMT5 is a methyl transferase that, among other proteins, targets several spliceosome proteins. Its deletion has been proved to provoke major splicing alterations [25, 26]. We analyzed two independent RNAseq assays that were conducted at different times probing the same biology. Experiments *A* (GSE149429) and *B* (GSE149430) were originally carried out to analyze splicing alterations in the PRMT5 knockout mutant in *Arabidopsis thaliana*. Both assays probed the PRMT5-KO and wild-type transcriptomes in Columbia ecotype plants as part of larger and different studies (see Material and Methods 3.4).

The rationale behind our analysis was two-fold. For one hand we wanted to assess for ASpli2 detection performance in a more realistic setup. For the other we wanted to take advantage of these datasets to quantitatively estimate the reproducibility of discoveries, i.e. we wanted to explore the consistency and robustness of experimentally identified alternatively splicing events in biologically related systems.

#### 2.3.1. Reproducibility assessment

We analyzed RNAseq assays *A* and *B* with ASpli2 and the other considered algorithms. For ASpli2, we used the same detection-call criteria specified in Section 2.2. Default parameters were considered to run the other tested methodologies (command lines used to execute them were included as Supplementary Material 8.2). For LeafCutter and rMATS we considered events presenting fdr corrected pvalues smaller than 0.05 and changes in junction inclussion indices larger than 0.1. For MAJIQ we sought for events presenting a posterior probability larger than 0.95, of having a change in inclussion index larger than a 0.2 level. Overall, 6350, 951, 412 and 158 genomic regions affected by altered splicing patterns were reported by ASpli2, LeafCutter, MAJIQ and rMATS algorithms respectively in at least one experiment.

In Table 3 we summarized reproducibility statistics for each examined methodology (a more in-depth comparison of discoveries was included as supplementary material in Section 8.5). Column *universe* of Table 3 reports the actual number of sub-genic regions that, upon passing different pre-filtering steps, were actually examined for statistically significant changes in splicing patterns. The extent of this background list was noticeably larger for ASpli2 as our methodology tested not only junction-related signals but also alterations in the usage of genomic bins. Columns *A* and *B* outline the number of regions reported as differentially spliced in each experiment and column *A* ∩ *B*, the discovery intersection size (i.e. number of sub-genic regions reported as differentially spliced in both data-sets). In parenthesis, we included the *overlap coefficient value*, defined as *A* ∩ *B/* min(*A, B*). Expected overlaps, fold enrichment (i.e. ratio between observed and expected overlaps) and p-values were estimated using the SuperExactTest R-package [30] and reported in *EO, FE* and *pval* columns respectively.

**Table (3).** Reproducibility statistics. The numbers of statistically analyzed genes (after prefiltering steps) for each algorithms are shown in the *universe* column. The number of splicing events reported for each experiment and the number of concordant discoveries are displayed at columns *A, B*, and *A*∩*B* respectively. The expected overlap, fold enrichment level and significance pvalue are displayed in columns EO, FE and pval respectively

| Method | universe | A | B | $A \cap B$ | EO | FE | pval |
| --- | --- | --- | --- | --- | --- | --- | --- |
| ASpli2 | 140191 | 4687 | 3904 | 2241 (0.57) | 130.5 | 17.2 | 0.0e+00 |
| leafCutter | 8113 | 603 | 675 | 327 (0.54) | 50.2 | 6.5 | 3.6e-219 |
| MAJIQ | 16441 | 277 | 284 | 149 (0.54) | 4.8 | 31.1 | 9.5e-203 |
| rMATS | 2405 | 310 | 401 | 310 (1.00) | 19.4 | 16.0 | 0.0e+00 |

It can be seen from Table 3 that, for all the examined methods, the agreement between experiments was highly significant. In all cases, more than 50% of events detected in one experimental instance was also reported in the other. At the same time it can be appreciated that ASpli2 provided the largest (and highly significant) overlap-set. Noticeably, the total number of concordant splicing-affected genomic regions detected by ASpli2 presented up to a 15-fold increase with respect to the size of concordant sets reported by others methodologies.

Overall our analysis showed that results obtained at different and independent experimental instances were reproducible, in the sense that statistically significant agreement was found for every methodology. These results were robust against using different overlap quantification criteria (see Sup.Mat 8.5). In this matter, and similarly to the results obtained on the synthetic dataset, our results on PRMT5 data showed that ASpli2 displayed high recall levels providing the largest list of concordant discoveries between experiments.

#### 2.3.2. Data consolidation

Up to now, we focused on the analysis of the intersection of set of discoveries as a measure of coherence of the results. Now we wanted to illustrate how ASpli2 capabilities to deal with complex experimental designs can be used to integrate experimental results in a more statistically sound way.

ASpli2 was used to consolidate datasets A and B considering a simple generalized linear model:

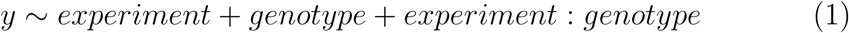

‘experiment’ was a fixed effect to cope with specific technical biases, and the ‘genotype’ factor was meant to capture the PRMT5 vs wild-type effect. The third term was an interaction term, and was used to enforced the exclusion of non-coherent signals between experiments.

ASpli2 detected 4360 genomic regions displaying strong evidence of a genotype effect (fdr *<* 0.05). In addition, 99% of these PRMT5-related events (4314 out of 4360) passed a filtering step to enforce they presented nodetectable evidence of experiment-genotype interactions (experiment:genotype associated fdr *>* 0.5). These 4314 events defined the consolidated AB data set.

We found that 99% (2209 out of 2241) of the concordant discoveries independently detected in both assays were also included in the consolidated dataset (we included a Venn diagram of the discoveries reported for experiments A, B, and the consolidated data-set AB in Sup.Fig. S7). Noticeably, the consideration of the AB data-set allowed to almost double the number of detected genomic regions displaying robust evidence of differential splicing patterns.

#### 2.3.3. PRMT5 RT-PCR detected events

The PRMT5 methyltransferase has been the target of many studies as deficiencies in this protein causes genomewide splicing alterations[26, 27, 28]. In this section we focused on two specific works that provided independent RT-PCR validated lists of splicing alterations linked to PRMT5 in Arabidopsis thaliana [25, 28].

For one hand, Deng and collaborators studied PRMT5 mutant *Arabidospis thaliana* plants and presented a list of 12 RT-PCR validated intron retention events (see Fig 2 in [28]). On the other, using the same biological model, Sanchez and collaborators indentified changes in alternative splicing using a high-resolution qRTPCR panel that included several known alternative splicing events [31]. They found that PRMT5 mutants had significant alterations in 44 events which included exon skipping, alternative donor and acceptor splice sites, as well as intron retention events (Supplementary Table 4 in [25]).

**Table (4).**

| Method |  | Deng 2010 | Sanchez 2010 |
| --- | --- | --- | --- |
| RT-PCR |  | 12 | 44 |
| ASpli2 | AB | 8 [8,4] | 23 [21,13] |
|  | A | 10 [8,5] | 24 [19,7] |
|  | B | 9 [8,2] | 20 [18,6] |
| LeafCutter | A | 3 | 16 |
|  | B | 3 | 17 |
| MAJIQ | A | 5 | 8 |
|  | B | 2 | 8 |
| rMATS | A | 1 | 12 |
|  | B | 1 | 3 |

We aimed to contrast these findings with the results reported by the different methodologies on datasets A and B. In Table 4 we summarized, for each study, the number of concordant findings uncovered by different algorithms on datasets A and B. Quantities between brackets represent the number of ASpli2 discoveries reported by coverage and junction-based signals respectively. It can be seen that ASpli2 recovered the largest number of events and that the majority of ASpli2 validated discoveries originated in differential coverage signal calls. Had we only considered junction related detection-signals, ASpli2 would have achieved similar levels of agreement than the other junction-based algorithms (for instance we got a similar performance than LeafCutter on Sanchez data-set for the consolidated case).

In Table S2, included as supplementary material, we further characterized the agreement between the 23 splicing events that ASpli2 uncovered for the consolidated AB case, and Sanchez qRT-PCR validated events. It can be seen that in 15 out of the 23 cases (65%), the very genomic region probed by the PCR analysis was recognized by ASpli2. For the other 8 cases, AS-pli2 detected actually occurrying changes in isoform usage but from splicing signals originating at genomic locations not probed by the PCR primers (See Supplementary Figures - TODO: ACA VAN SAHIMI PLOT DE EVENTOS PCR).

### 2.4. ASpli2 scalability analysis

In this section we leveraged on a mid-size RNAseq study presented by Ren and collabrators to characterize aberrant splicing patterns occurying in prostate cancer patients [29]. We aimed to analyze this sample-paired assay to see how ASpli2 performance (statistical power, precision, time and memory requirements) scaled with the number of samples. In particular, we followed the approach suggested in [32] to characterize ASpli2 in terms of statistical power and expected false discovey rate for a varying number of samples.

#### 2.4.1. Statistical power

Ren and collaborators presented a comprehensive study of splicing alterations detected using RNAseq transcriptome profiles of 14 primary prostate cancers and their paired normal counterparts from the Chinese population [29]. On average, the 28 fastq files presented 34.6 *±* 1.7 million reads per sample and 31.4 *±* 1.6 millions of them were actually mapped to the EN-SEMBL HG38.98 version of the human genome (see Material and Methods 3.6). The genome’s GTF and BAMs files were then used as inputs to drive an alternative splicing paired-sample analysis with ASpli2. We considered the following model to identify genomic regions differentially spliced in tumor samples compared to normal tissue controls:

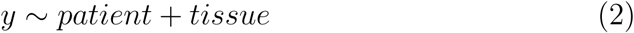

The ‘patient’ term served to pair tumor and normal tissue samples coming from the same individual. The two-level ‘tissue’ factor reported average differences between tumor and normal cases over the observed population of patients.

In order to study the dependency of the statistical power on the number of samples, we sampled without replacement (10 times) subsets of 3, 5, 7 and 10 individuals. For each case, we reported, in the first column of Table 5, the median (and standard error, in brackets) of the number of genomic regions found to be alternatively spliced between tumor and normal samples.

**Table (5).**
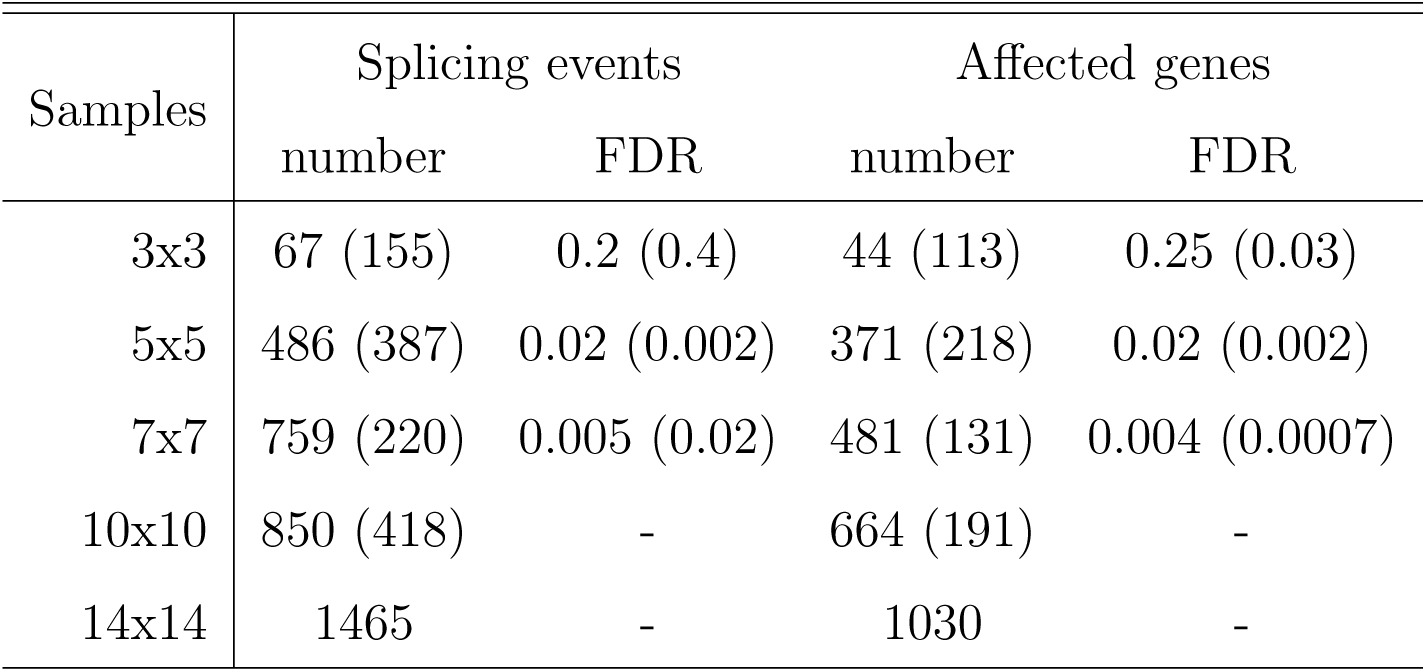
Summary of the 10-fold bootstrapped analysis of ASpli2 performance on the prostate cancer data set. For each number of paired samples (first column) the median number of genomic-regions displaying a statistically significant ‘tissue’ effect were included in the second column. Median values of false discovery rate estimations obtained from the analysis of normal-tissue samples were shown in the third column. Standard error estimation were reported between brackets.

In order to estimate false discovery rates we considered mock comparisons between normal samples (we sampled 10 times normal tissue samples of 3vs3, 5vs5 and 7vs7 individuals). We then estimated FDR as the ratio between the number of mock discoveries and the median number of discoveries found in true comparisons of the same number of samples. In the second column of Table 5 median and standard errors (in brackets) were reported.

It can be seen from Table 5 that the median number of detected splicing events increased with the number of examined samples, up to a maximum of 1465 events obtained when the 28 paired samples were considered. The large variability observed between bootstrap realizations was consistent with the large variability already observed across prostate cancer transcriptomes (see [29] and Supplementary Material 8.7). FDR estimated values showed a huge decrease with increasing number of samples, and for the 5×5 case seemed to have already leveled off. Similar trends were observed when splicing alter-

ations were reported at the level of hosting genes (data not shown).

#### 2.4.2. Time and memory requirements

In Table 6 we reported median values and standard errors for the elapsed time and peak memory usage required for calculations (performed on single thread on an Intel Xeon Silver 4116 2.1GHz Lenovo ThinkSystem SR650)

**Table (6).** Summary of the 10-fold bootstrapped analysis of ASpli2 performance on the prostate cancer data set. For each number of paired samples (first column) the median number of genomic-regions displaying a statistically significant ‘tissue’ effect were included in the second column. Median values of false discovery rate estimations obtained from the analysis of normal-tissue samples were shown in the third column. Median time and memory used in the analysis were reported in the last two columns. Standard error estimation were reported between brackets.

| Samples | time [min] | memory peak[Gb] |
| --- | --- | --- |
| 3x3 | 67 (1) | 20.25 (0.38) |
| 5x5 | 111 (2) | 22.15 (0.20) |
| 7x7 | 156 (4) | 24.13 (0.03) |
| 10x10 | 231 (5) | 26.57 (0.04) |
| 14x14 | 348 | 30.07 |

Execution time scaled linearly with the number of paired samples at a rate of 25.5 minutes per pair of samples (about 90% of execution time was used for BAMs reading and feature counting). The memory peak column shows that RAM requirement linearly scaled with the number of samples at a rate of about 880*Mb* per sample pair. A simple extrapolation suggests that about 65Gb should be enough to handle 100 samples of the same sequencing

depth (∼ 3.510^6^ reads per sample).

## 3. Material and Methods

### 3.1. Differential analysis scheme

ASpli2 leverages on the statistical framework developed by Smyth and collaborators, implemented in the edgeR R-package [24, 20], to assess for statistically significant changes in gene-expression, bin coverage and junction splicing signals. Under this approach, count data is modeled using a negative binomial model, and an empirical Bayes procedure is considered to moderate the degree of overdispersion across units.

*Differential expression signals*. Differential expression signals are estimated via generalized linear models (GLM). This approach allows ASpli2 to deal with complex experimental designs, i.e. contrasts can be tested in experiments with multiple experimental factors. Using this statistical setting, for each gene, ASpli2 quantifies differential gene expression signals reporting the corresponding log-fold change, p-value, and FDR adjusted q-values.

*Differential splicing signals*. In order to study splicing patterns, gene expression changes should be deconvolved from overall count data. On a very general setting, what we are looking for is to test whether a given unit of a certain group of elements displays differential changes respect to the collective or average behavior. ASpli2 uses this general idea to assess for statistically significant changes in splicing patterns probed with different genomic features:

- bin-coverage signal: ASpli2 assesses for differential usage of bins comparing bin’s log-fold-changes with the overall log-fold-change of the corresponding gene.
- junction anchorage signal: For every experimentally detected junction, ASpli2 analyzes differential intron retention changes by considering log-fold-changes of a given experimental junction relative to changes in coverage of left and right junction flanking regions.
- junction locale signal: In the same spirit than MAJIQ and LeafCutter, ASpli2 defines junction-clusters as sets of junctions that share at least one end with another junction of the same cluster (see Panel E of Figure S8). In order to characterize changes for a given junction along experimental conditions, ASpli2 weighs log-fold-change of the junction of interest relative to the mean log-fold-change of junctions belonging to the same cluster.

ASpli2 makes use of the functionality implemented in the diffSpliceDGE function of the edgeR package to perform all of this comparisons within a unified statistical framework. Given a set of elements (i.e. bins or junctions) of a certain group (i.e. genes, anchorage group or junction-cluster), a negative binomial generalized log-linear model is fit at the element level, considering an offset term that accounts for library normalization and collective changes. Differential usage is assessed testing coefficients of the GLM. At the single element-level, the relative log-fold-change is reported along with the associated p-value and FDR adjusted q-values. In addition a group-level test is considered to check for differences in the usage of any element of the group between experimental conditions (see *diffSpliceDGE* documentation included in edgeR package for details [24]).

### 3.2. Filtering and detection criteria

Statistical analysis of differential splicing is performed only on expressed genes (i.e. read counts spanning the gene genomic range should be larger than a minimal number of reads, 5 by default, across all the samples of the contrasted conditions). Furthermore, analyzed bins and junctions should present a minimal number of counts (5 by default) in every replicate of at least one contrasted condition. Additionally, marginally present junctions are filter-out looking at the maximal value of their *participation* coefficient, defined as the relative abundance of a given junction within its group for a given experimental condition.

Besides statistical figures of merit, ASpli2 provides additional statistics and parameters in order to ease the identification of biologically relevant events. For instance, magnitude of change in inclusion or strength indices (see Table S1) between experimental conditions, are also reported in order to filter-out weak events. In this way, a bin is called differentially-used by ASpli2 if it displays statistically significant coverage changes (fdr *<* 0.05, by default) and, additionally, one of the two supplementary conditions hold: either the bin fold-change level is greater than a given threshold (3 fold changes, by default) or changes in inclusion levels of bin-supporting junctions (Δ*PIR* or Δ*PSI* according to the bin class, see Table S1) surpasses a predefined threshold (0.2 by default).

Anchorage splicing signals, on the other hand, are reported whenever statistically significant changes are found at the cluster level (cluster.fdr *<* 0.05 by default) for the considered {*J*_1_, *J*_2_, *J*_3_} junction set (see upper panel of Fig S8-D) and, at the same time, 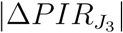 is larger than a given threshold (0.3 by default).

Finally, junction locale differential splicing signals are reported whenever statistically significant changes are found at the cluster level (cluster.fdr *<* 0.01 by default) for the analysed junction cluster {*J*_1_, …, *J*_*S*_, …, *J*_*n*_} (see S8-E) and, at the same time, there is at least one junction *J*_*S*_ within the cluster presenting statistically significant changes at the single unit level (junction.fdr *<* 0.05, by default) with 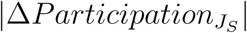 larger than a given threshold (0.3 by default). In the case that statistically significative changes were detected at the unit-level for more than one junction of a given cluster, the one displaying the largest participation change was considered and reported as the cluster’s representative junction.

### 3.3. Splicing simulation

We implemented a computational pipeline relying on the Flux Simulator (FS) software [33] in order to produce a controlled set of splicing events. We first used FS to generate a transcript abundance distribution template to spread 15 *×* 10^6^ molecules among the 10646 available transcript variants of the 8433 genes of chromosome-one of the *Arabidospis thaliana* genome. Then, we generated a ‘treatment’ set of samples altering the original molecule distribution in order to simulate genome-wide differential changes in gene expression and splicing patterns.

Finally we simulated biological replicates from these two *seed* transcriptomes, considering a Gamma distribution for molecule abundances to build ‘control’ and ‘treatment’ sample sets. We chose to work with a *CV* = 0.1 level of variability in gene abundance between replicates. Therefore, we considered *shape* (*k* = 100), and *scale* (*θ* = 0.01*µ*) parameter values, where *µ* was the gene expression level in the corresponding *seed* transcriptome used for replicate generation.

Simulated changes in variant concentrations produce patterns of differential usage at bin and junction levels according to the exonic architecture of the different gene variants. For instance, a splicing alteration that involves switching between Isoform 1 and Isoform 3 of the gene depicted in Figure S8-(A) is expected to produce differential usage signals for the first and third exonic bins. In our case we simulated changes in variants usage for 915 genes that should altered, in principle, the coverage signal of 2451 bins. It is worth mentioning that as alternative splicing was modeled exclusively through differential variant usage, no intron retention events were simulated in the synthetic data set.

Several examples of splicing simulated events are depicted in Sup. Figs S4,S6,S5. Examples, scripts and additional material to reproduce the ASpli2 analysis over this dataset can be found at the gitlab repo: https://gitlab.com/ChernoLab/aspli2_sm.

### 3.4. PRMT5 datasets

The goal of these studies was to compare the transcriptional profile (RNA-seq) of wild type and PRMT5 Arabidopsis mutants plants grown under continuous light at 22 degrees centigrades.

Dataset A (GSE149429): WT (Col) and PRMT5 mutants seeds were grown on Murashige and Skoog medium containing 0.8% agarose, stratified for 4 d in the dark at 4 C, and then grown for fifteen days under continuous white light at 22C Whole plants were harvested after 15 d. Total RNA was extracted with RNeasy Plant Mini Kit (QIAGEN) following the manufacturers protocols. To estimate the concentration and quality of samples, NanoDrop 2000c (Thermo Scientific) and gel electrophoresis were used, respectively. Libraries were prepared following the TruSeq RNA Sample Preparation Guide (Illumina). Briefly, 3 g of total RNA was polyA-purified and fragmented, and first-strand cDNA synthesized by reverse transcriptase (SuperScript II; Invitrogen) and random hexamers. This was followed by RNA degradation and second-strand cDNA synthesis. End repair process and addition of a single A nucleotide to the 3 ends allowed ligation of multiple indexing adapters. Then, an enrichment step of 12 cycles of PCR was performed. Library validation included size and purity assessment with the Agilent 2100 Bioanalyzer and the Agilent DNA1000 kit (Agilent Technologies)

Dataset B (GSE149429): WT (Col accession) and PRMT5 mutant plants were grown for nine days under continuous white light at 22 degrees centigrades or exposed for 1 or 24 h to 10C on the 9th day, before harvesting. Then the transcriptional profile of these plants was analyzed using RNA-seq. WT (Col) and PRMT5 mutants seeds were grown on Murashige and Skoog medium containing 0.8% agarose, stratified for 4 d in the dark at 4 C, and then grown for nine days under continuous white light at 22C. Whole plants were harvested after 9 d. Total RNA was extracted with RNeasy Plant Mini Kit (QIAGEN) following the manufacturers protocols. To estimate the concentration and quality of samples, NanoDrop 2000c (Thermo Scientific) and gel electrophoresis were used, respectively. Libraries were prepared following the TruSeq RNA Sample Preparation Guide (Illumina). Briefly, 3 g of total RNA was polyA-purified and fragmented, and first-strand cDNA synthesized by reverse transcriptase (SuperScript II; Invitrogen) and random hexamers. This was followed by RNA degradation and second-strand cDNA synthesis. End repair process and addition of a single A nucleotide to the 3 ends allowed ligation of multiple indexing adapters. Then, an enrichment step of 12 cycles of PCR was performed. Library validation included size and purity assessment with the Agilent 2100 Bioanalyzer and the Agilent DNA1000 kit (Agilent Technologies).

On average, 19.3 *±* 5.3 million 100 long and 28.3 *±* 7.7 million 150 long paired-end reads were generated per sample library for datasets *A* and *B* respectively. For both cases more than 96% of reads were uniquely mapped to TAIR10 Arapidopsis genome using STAR (command-line invocation was included in Sup Mat 8.2).

### 3.5. Overlap analysis

We followed the procedure outlined in Supplementary Material 8.3 to map events reported by each of the considered method to a common set of genomic coordinates. Overlaps were then estimated using the *findOverlaps* function of the *IRanges* package of R [34].

### 3.6. Prostate cancer dataset

Fifty-six paired fastq files from the E-MTAB-567 experiment were downloaded from the ArrayExpress server. Reads were aligned against ENSEMBL HG38.98 reference genome using the STAR aligner with default parameters and a junction overhang=89.

### 3.7. Code availability

ASpli2 package is freely available at https://gitlab.com/ChernoLab/aspli2, and will be part of the next Bioconductor release (October 2020). Examples, scripts and additional material to reproduce our analysis can be found at the gitlab repo: https://gitlab.com/ChernoLab/aspli2_sm.

## 4. Discussion

RNA high-throughput sequencing methods provide powerful means to study alternative splicing under multiple conditions in a genome-wide manner. However, the detection and understanding of general splicing patterns still present considerable technical challenges. Here we presented ASpli2, a computational suite to comprehensively test bin coverage and junction usage differential splicing signals.

The analysis methodology implemented in ASpli2 came out as a result of several software maturation cycles of our in-house splicing analysis procedures. Over the last years, the presented core functionality has been extensively used in different projects to study: the role of AS in circadian rhythms and light response [35, 36, 37, 38, 39] as well as AS in spliceosome mutants [40, 41] in A.thaliana model organism. In addition, ASpli2 in-house versions have been used to study AS and rhythmic behavior in D.melanogaster [42] and to characterize AS in dengue’s viral infection in humans [43].

In order to quantify ASpli2’s performance we compared it against three different state-of-the-art methodologies: LeafCutter [23], MAJIQ [44] and rMATS [22]. As a general rule we considered default parameters to run these analysis pipelines for our intention was not to present here an extensive benchmark between bioinformatics approaches, nor to propose the definitive analysis methodology. Rather we wanted to establish whether ASpli2 produced reasonable and competitive results.

Different scenarios were considered to chart ASpli2 performance. We first analyzed a synthetic data-set and quantified the ability of each considered methodology to detect splicing changes in terms of precision and sensitivity figures of merit. Using this controlled dataset we found that all the analysed methods presented rather high precision levels. However ASpli2 systematically displayed larger recall values (∼ 40%), mainly because the use of coverage signals (see Fig 1). This is an important result as highlights the benefits of not loosing effective sequencing depth by relaying not only on junction information but on the complete set of reads of RNA-seq runs.

We then aimed to outline ASpli2’s performance over more realistic setups. As no internal gold-standards are usually available for real world datasets we focused on the analysis of two independent RNAseq assays that probed the same biological conditions. This allowed us to quantify the consistence and coherence of outcomes produced by each methodology in terms of reproducibility of discoveries. Our results suggested that detection agreement between studies was highly significative for every methodology. However ASpli2 was far superior in terms of total number of concordant discoveries reported.

It is worth noting that a necessary condition implicit in this analysis was that biological variability largely exceeded possible technical biases between studies. Using ASpli2, we were able to consider a generalized linear model to define a consolidated dataset integrating data from both studies and verified that this was actually the case (Sec 2.4). In addition, the possibility to implement a two-factor model greatly improved the statistical power to uncover consistent discoveries. We could identify 4314 events displaying a statistically significative genotype effect and no evidence of experiment-genotype interactions. This represented almost a two-fold increase in the number of reproducible discoveries when compared against the naive integrative approach that merely considered the 2241 splicing events simultaneously detected in both studies.

An important aspect of the presented approach is that ASpli2’s core functionality is implemented along user-friendly functions that produce selfcontained output results for each step of the analysis. This is an important feature from the user’s perspective. It provides the user valuable intermediate information eventually facilitating the integration of ASpli2 with other analysis pipelines.

## 5. Conclusions

In this paper we presented ASpli2, a computational suite to study alternative splicing events. It is implemented as a flexible R modular package that allows users to fulfill gene-expression and splicing analysis following a set of simple steps.

Noticeably, ASpli2 can handle complex experimental designs using a unified statistical framework to assess for differential usage of sub-genic features and junctions. By combining statistical information from exons, introns, and splice junctions ASpli2 can provide an integrative view of splicing landscapes that might include canonical and non-canonical splicing patterns occurring in annotated as well as in novel splicing variants.

## Acknowledgements

We thank Ruben Schlaen, Julieta Mateos and Andres Romanowski for helpful discussions

## 7. Supplementary Figures

**Figure (S1).**
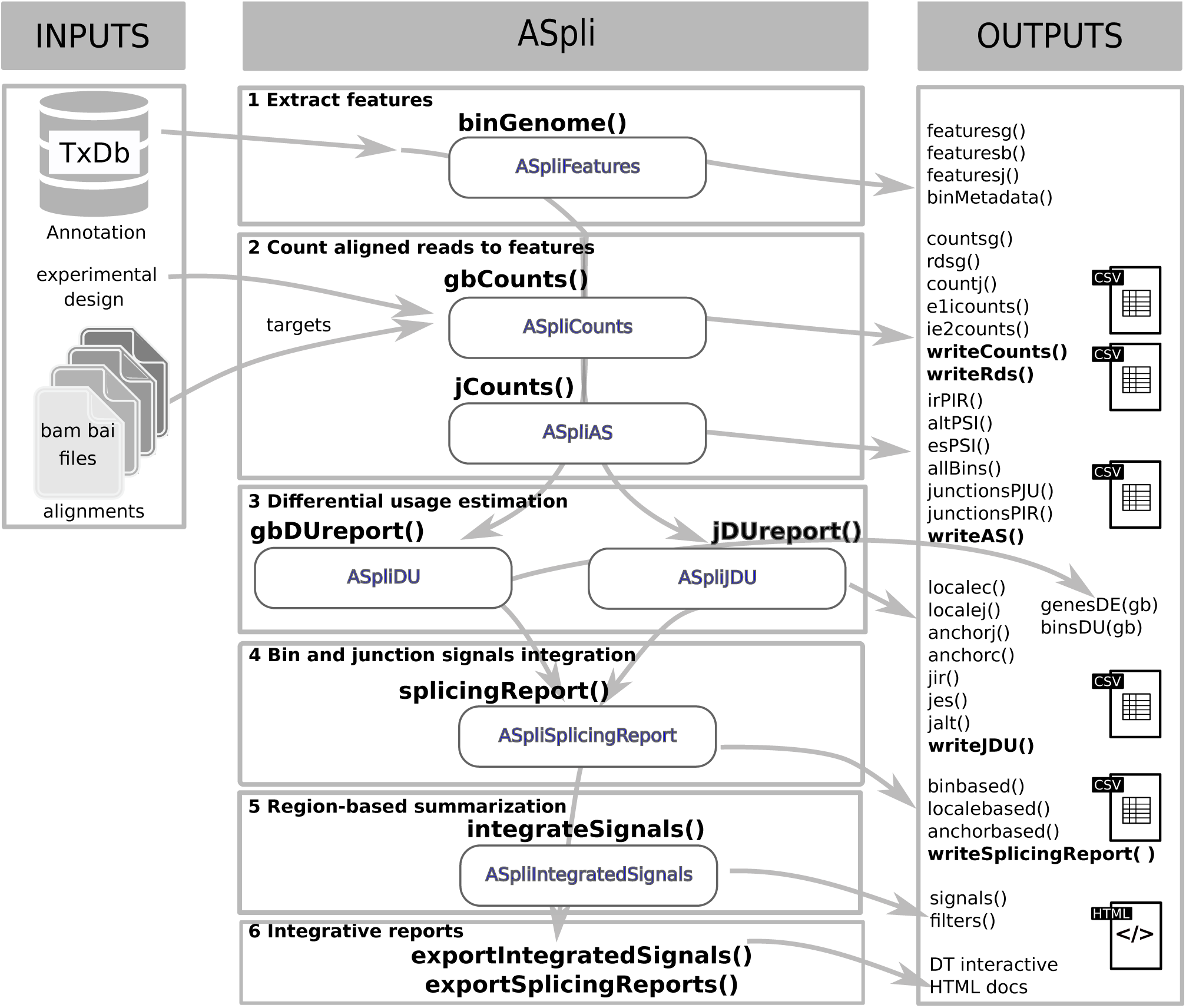
ASpli workflow. Rounded boxes are objects created by ASpli functions. Accesors and outputs are summarized in the right-most panel

**Figure (S2).**
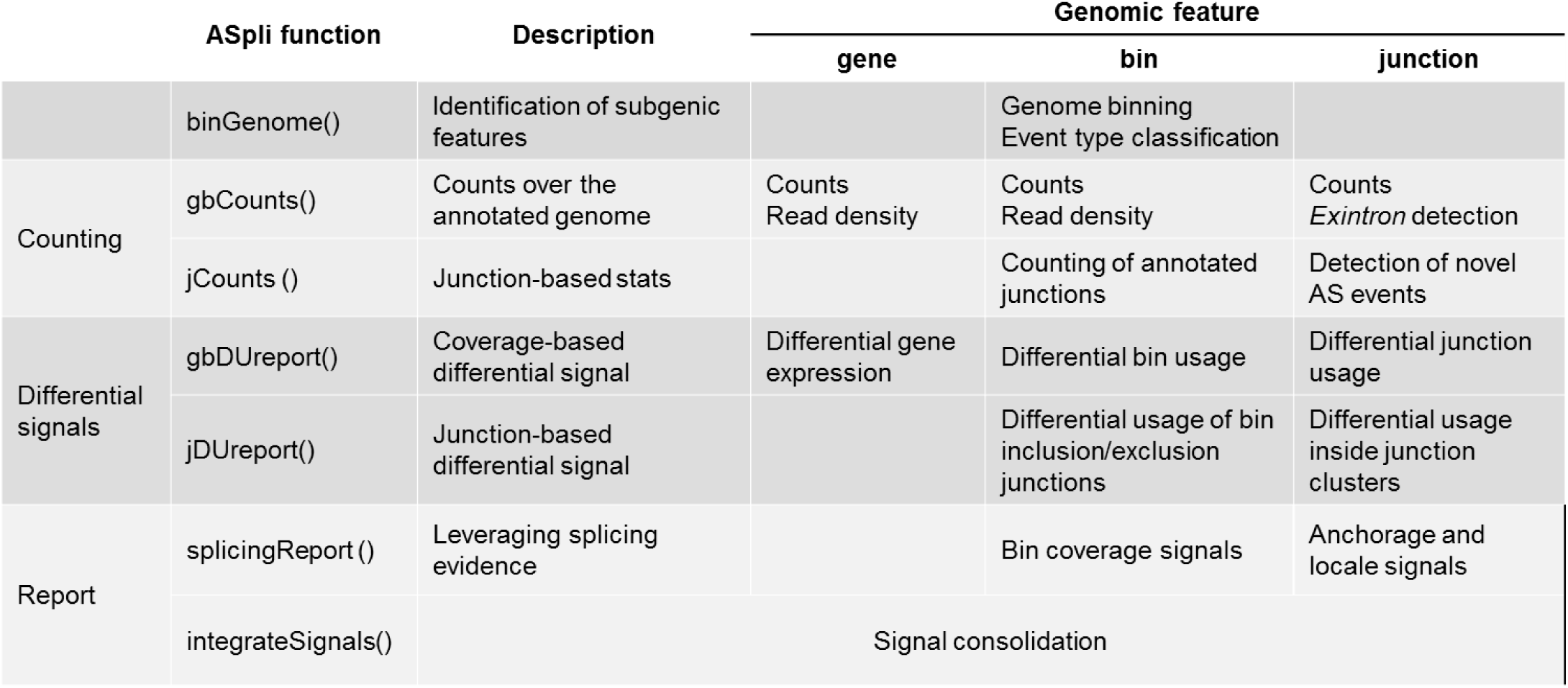
Summary of ASpli core functionality.

**Figure (S3).**
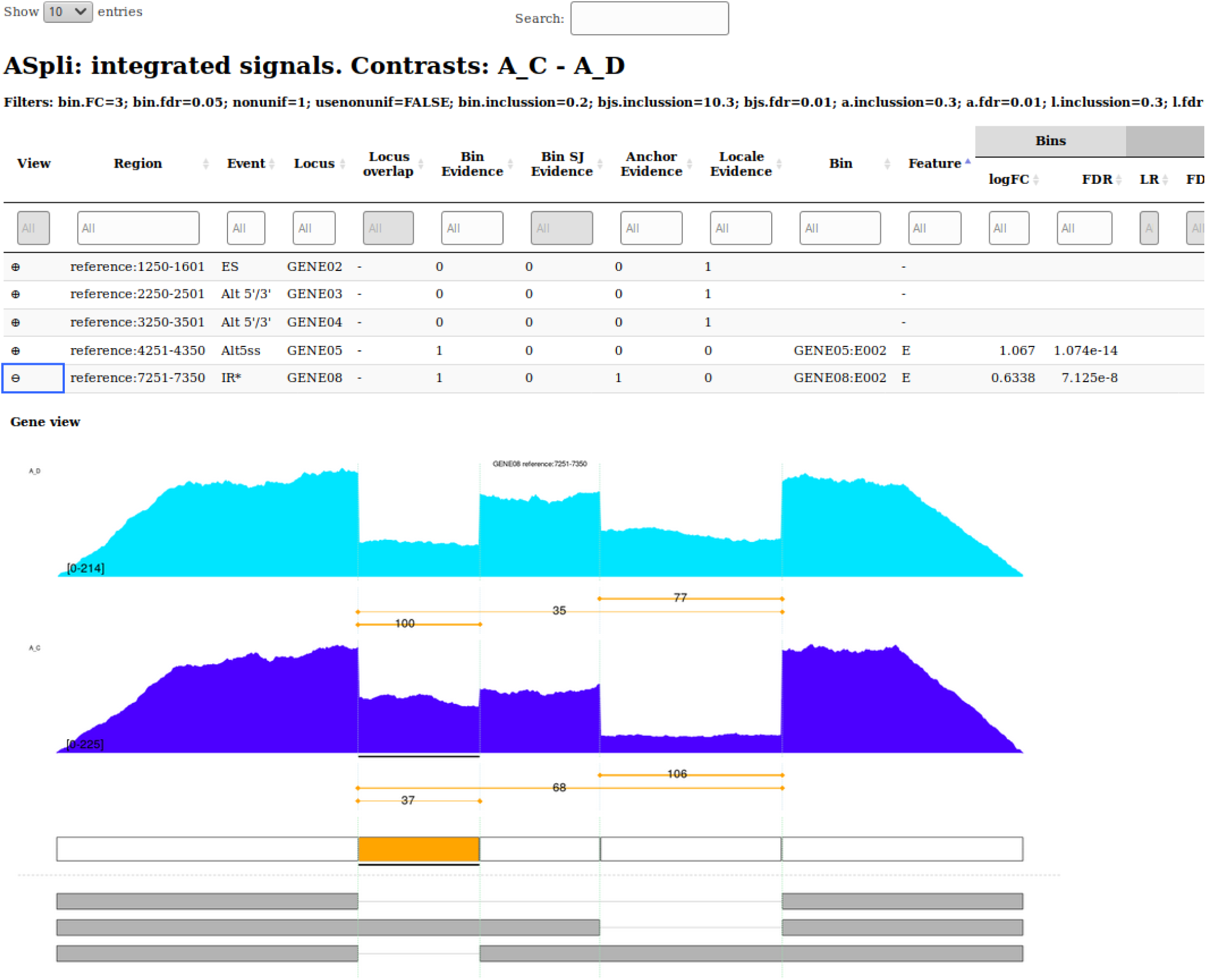
Example of DT html interactive report generated by *exportIntegratedSignals()* function

**Figure (S4).**
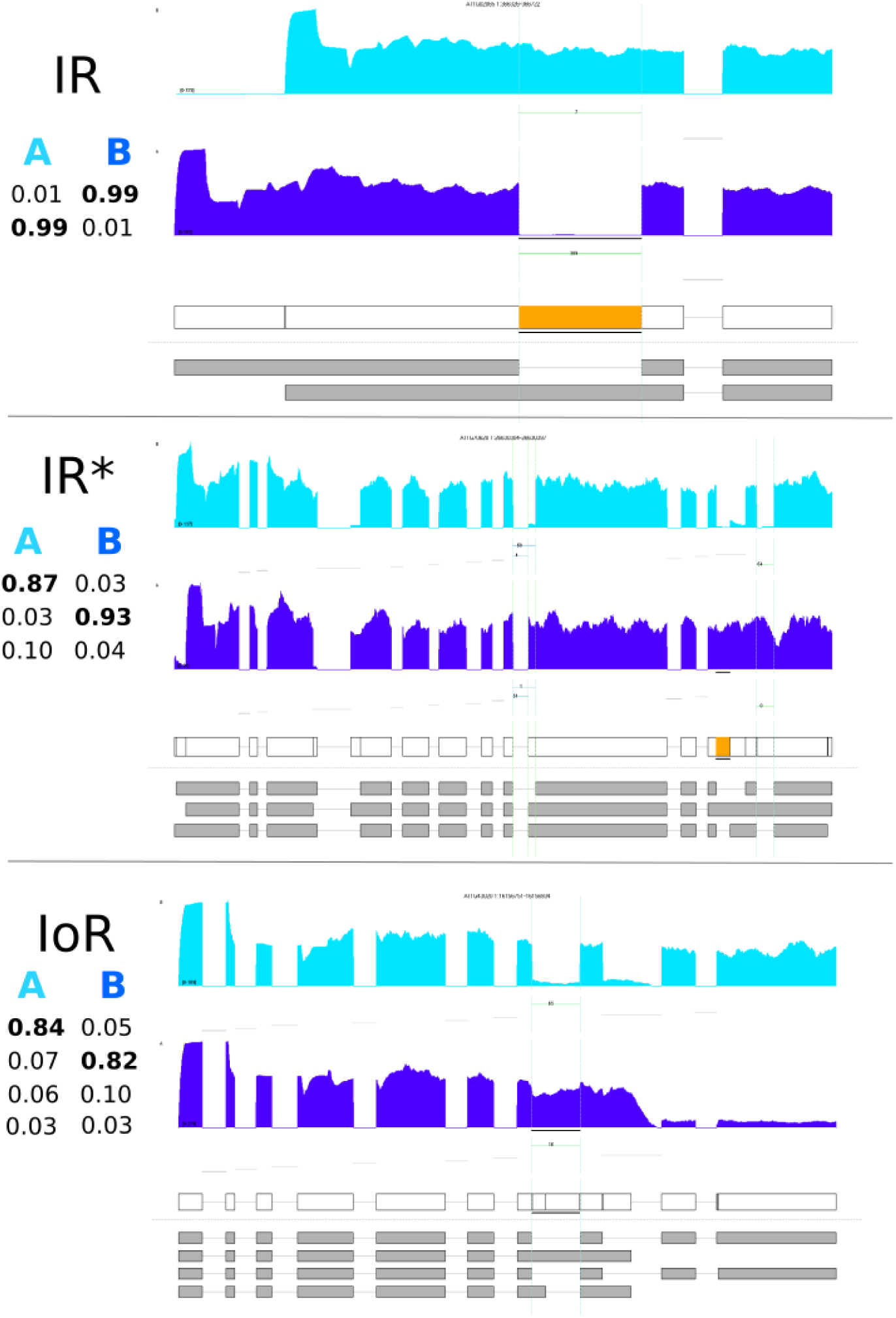
Examples of simulated IR-like splicing events. For each panel, the left layered table shows the relative concentration of each variant simulated for condition A and B. Orange boxes highlight the considered bin in each case.

**Figure (S5).**
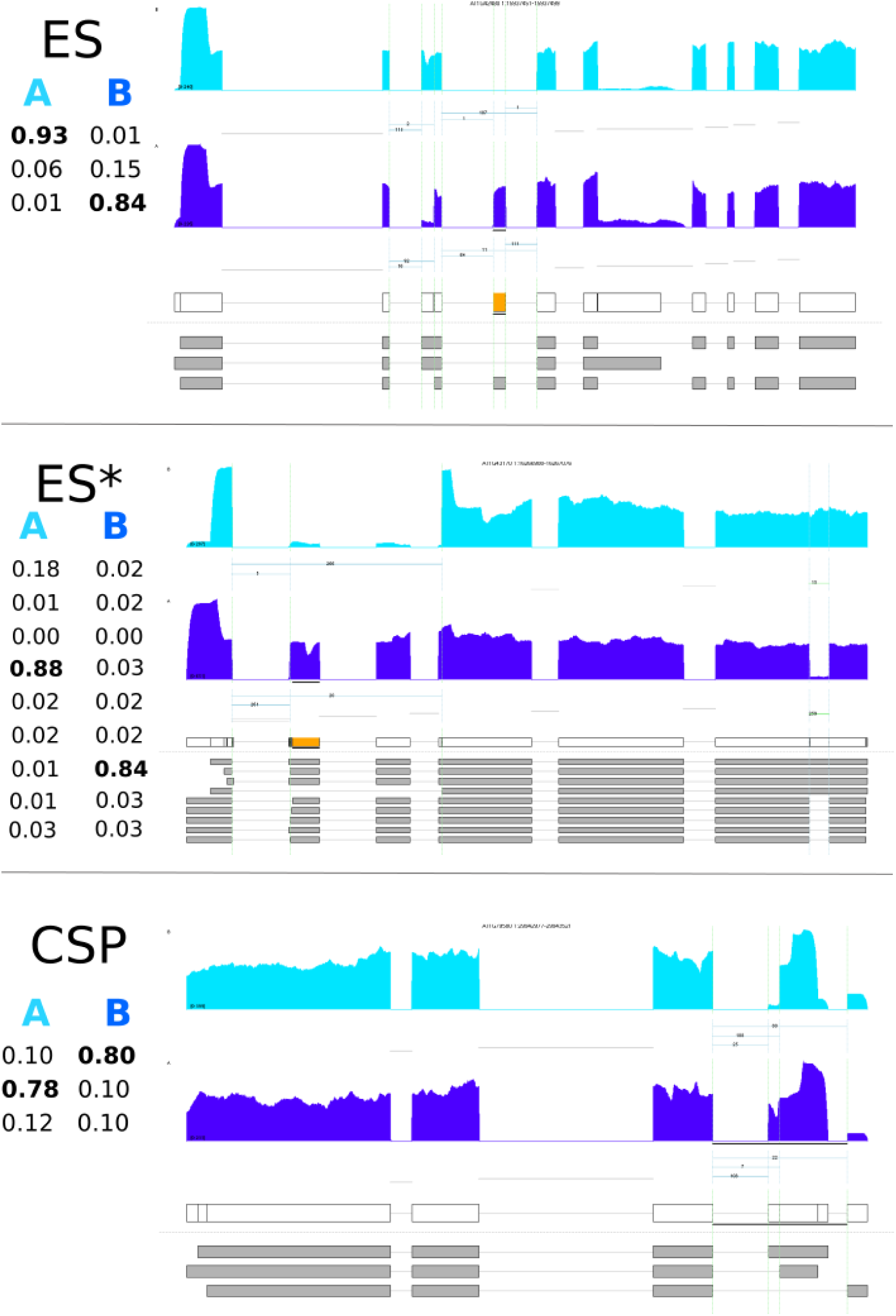
Examples of simulated ES-like splicing events. For each panel, the left layered table shows the relative concentration of each variant simulated for condition A and B. Orange boxes highlight the considered bin in each case.

**Figure (S6).**
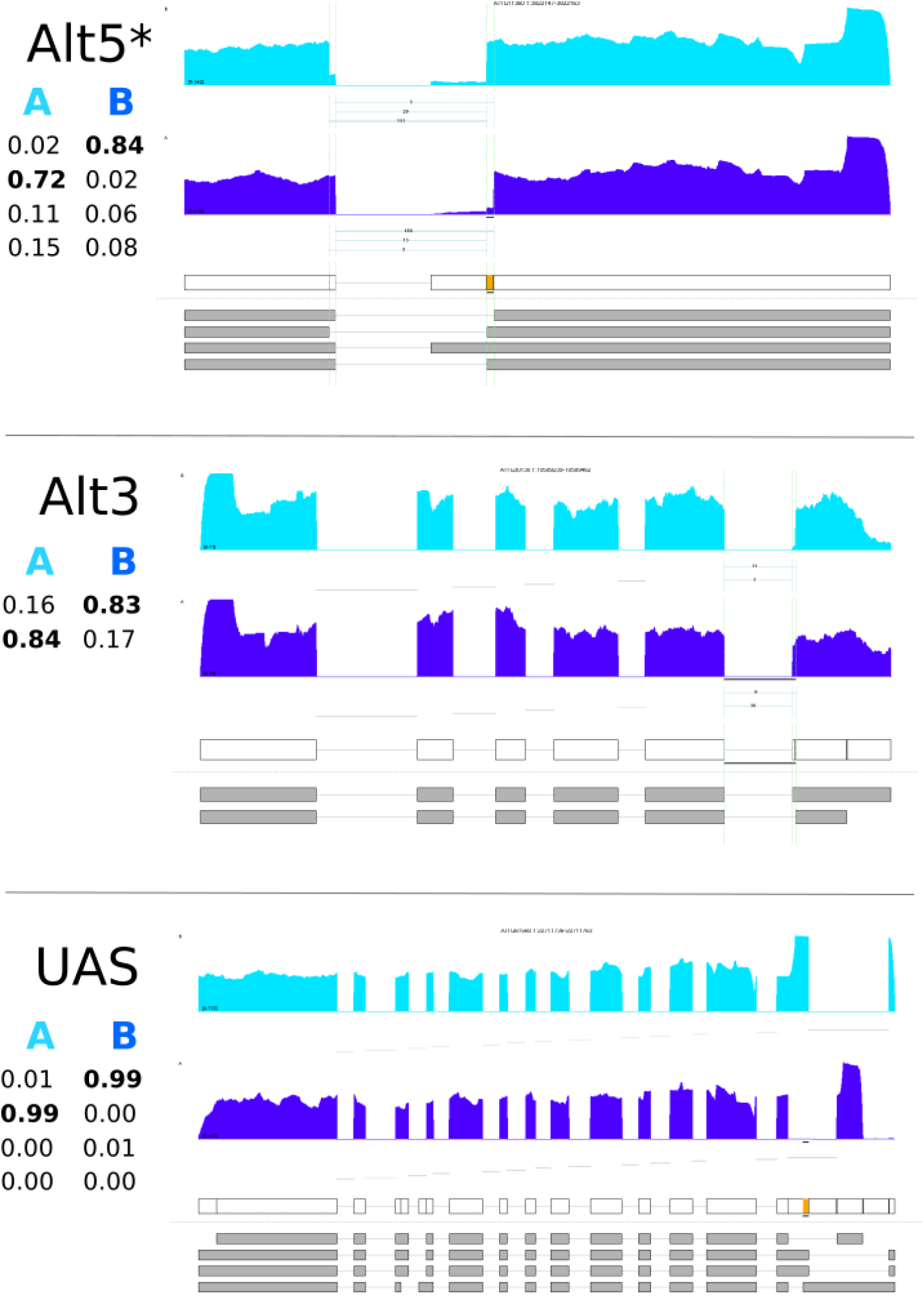
Examples of simulated Alternative start/end splicing events. For each panel, the left layered table shows the relative concentration of each variant simulated for condition A and B. Orange boxes highlight the considered bin in each case.

**Figure (S7).**
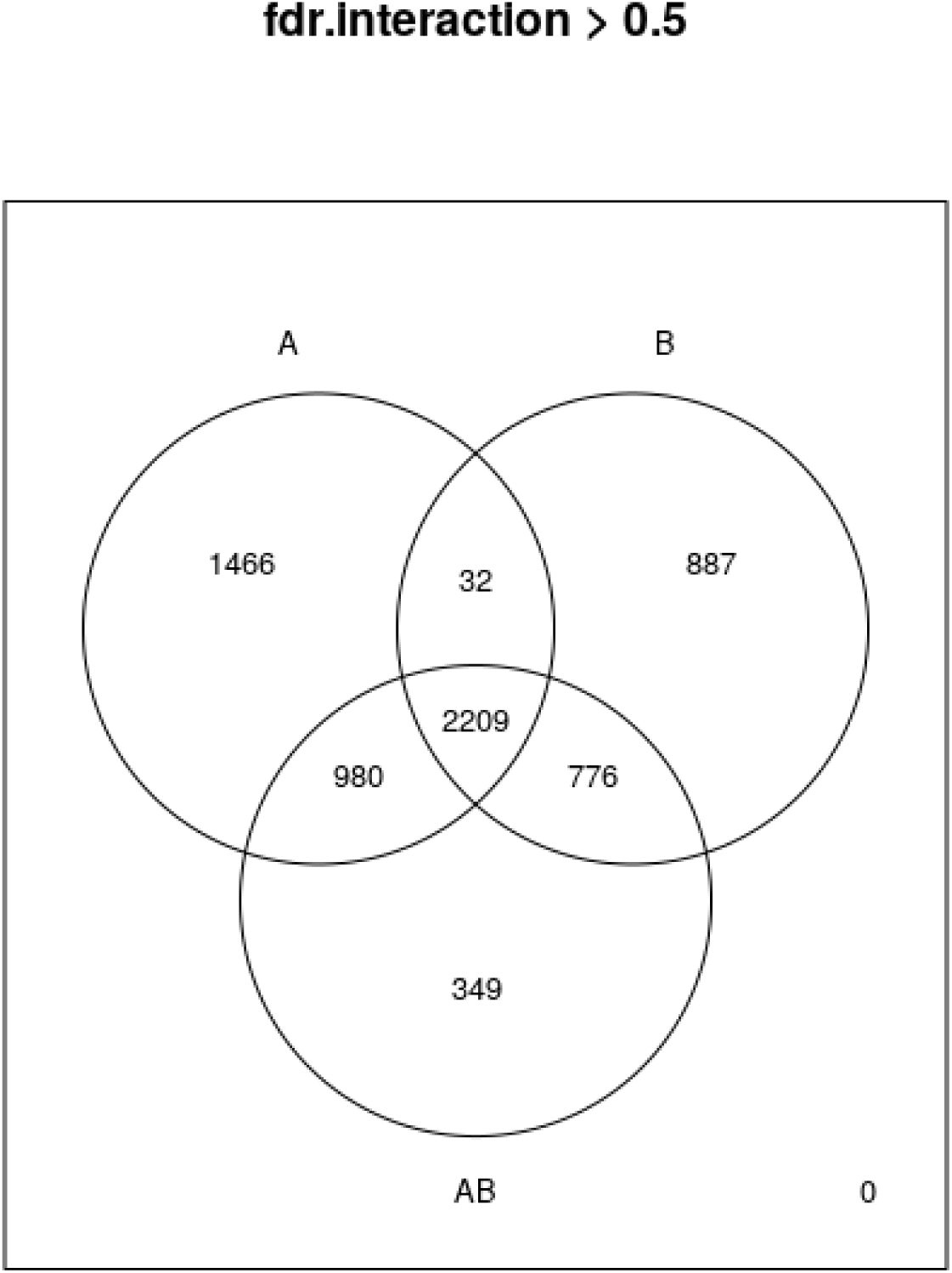
Venn diagram of alternative splicing events detected in experiments A, B, and the consolidated data set AB (i.e. events displaying strong evidence of a genotype effect (fdr*<* 0.05) and no-detectable evidence of experiment-genotype interaction (experiment:genotype associated fdr*>* 0.5)).

## 8. Supplementary Material

### 8.1. Feature counting in ASpli

#### 8.1.1. *Genomic feature extraction:* binGenome()

Sub-genic features are analyzed using user-provided annotation files. Exon and intron coordinates are extracted from annotation for multi-exonic genes. When more than one isoform exists, some exons and introns from different isoforms will generally overlap. In the same spirit of [18], exons and introns are then subdivided into non-overlapping sub-genic features dubbed bins, defined by the boundaries of different exons across transcript variants. In this way, these so defined *bins* are maximal sub-genic features entirely included or entirely excluded from any mature transcript.

Bins are flagged as: exonic (E), intronic (I) or alternative-splicing (AS) bins, depending on the exonic/intronic character of the bin across variants. In addition, original intronic (Io) bins are defined for every intronic region of annotated isoforms (see panel A of Figure S8).

As a general rule, the extreme portions of a transcript probed by RNAseq assays show a highly non-uniform coverage that might obscure differential usage analysis. ASpli flags bins that overlap with the beginning or ending of any transcript as *external*. An external bin of a transcript may overlap with a non-external one of another transcript. Whenever this happens the bin is still labelled as external. Additionally, in order to avoid confounding effects in the analysis of splicing events, ASpli identifies and flags loci where more than one gene is present in the genome.

*Local splicing classification model*. Each AS bin is further classified considering a three-bin *minimum local gene model*, that assigns splicing-event categories to a given bin based on the intronic/exonic character of the analyzed bin and its first neighbors (Figure S8, panel B).

For genes presenting two isoforms, this model is able to unambiguously assign a well defined splicing event to the analyzed bin: exon skipping (ES), intron retention (IR), alternative five prime splicing site (Alt5’SS), or alternative three prime splicing site (Alt3’SS) (see first row of panel B in Figure S8).

When more than two isoforms are present, we still found it useful to use the three-bin local model to segment follow up analysis. For these cases ASpli identify splicing events that involve: intronic subgenic regions surrounded by exons in at least one isoform (bin labelled as IR*), exonic subgenic regions surrounded by two introns in at least one isoform (bin labelled as ES*), exonic regions surrounded by intronic and exonic neighbor bins (bin labelled as Alt5’SS* or Alt3’SS*). When it is not possible to get a clear splicing-type assignation (see rows 2-5 of Figure S8), bins are labeled as *undefined AS* (UAS).

As a last step of the genomic feature extraction process, annotated junctions from all the transcripts are also identified. Junction coordinates are defined as the last position of the five prime exon (donor position) and the first position of the three prime exon (acceptor position).

#### 8.1.2. *Annotation based feature counting:* gbCounts()

Reads are overlaid on features derived from annotation, and count tables are produced at different genomic levels: genes, bins, and intron flanking regions used to identify and quantify intron retention events. Reads corresponding to annotated junctions are also tallied, along with genomic relevant information such as identity of spanned bins, and the existence of possible *exintronic* events [**?**].

**Figure (S8).**
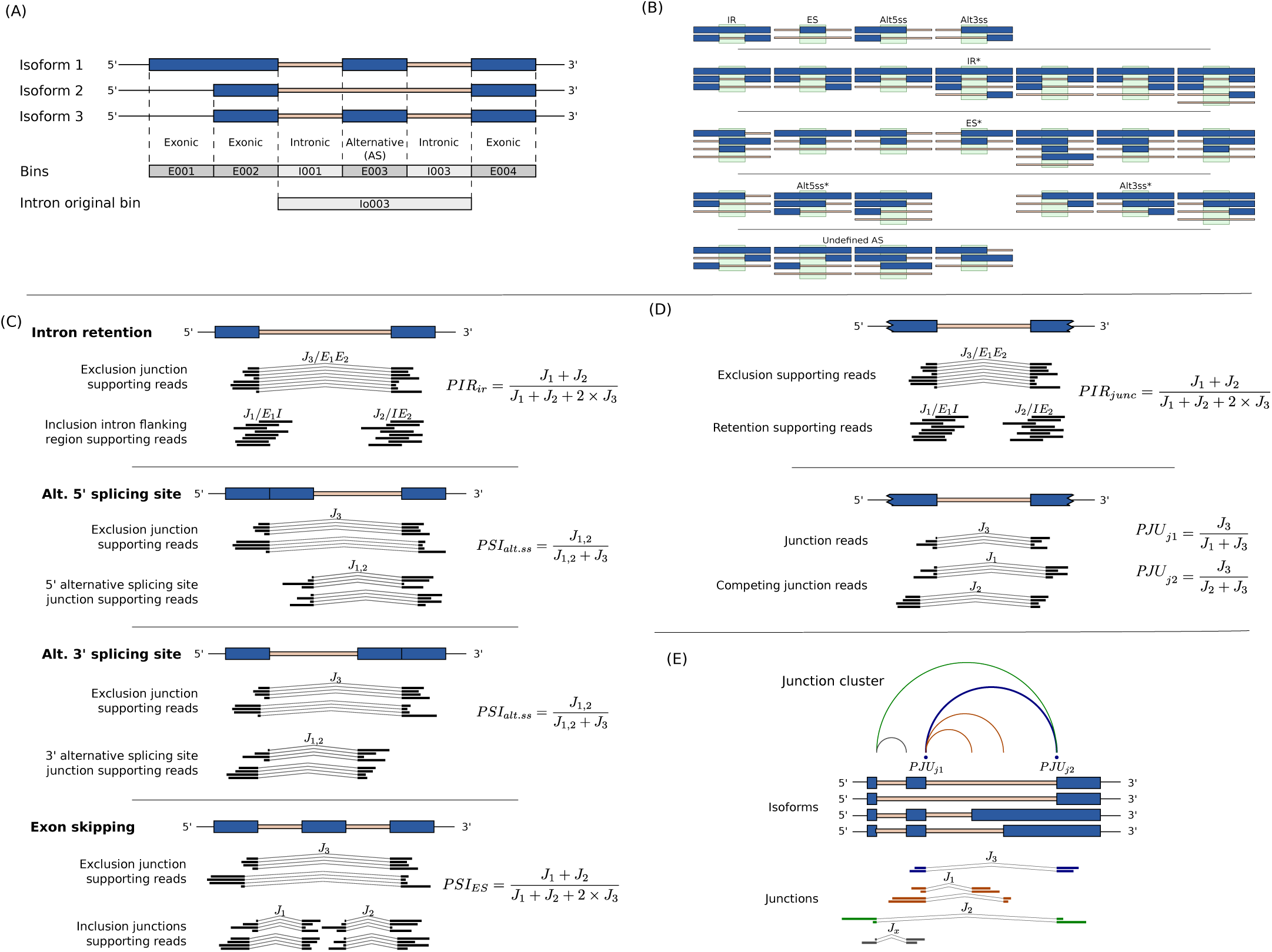
Panel (A) shows how bin-features are defined and classified as: exonic, intronic or intron original bins using genome annotation. The local splicing classification scheme is illustrated in panel (B). The definition of PSI and PIR metrics for bin features are pictured in panel (C). Definition of junction PIR and PJU statistics are shown in panel (D). Panel (E) shows a possible junction cluster and highlights the definition of type *J*_1_, *J*_2_ and *J*_3_ junctions for the analysis of PJU statistics for the blue junction.

#### 8.1.3. *De-novo junction counting:* jCounts()

ASpli takes advantage of experimentally detected splice junctions to perform two different type of analysis. For one hand, junction data is considered in order to provide junction support to AS events detected through bin coverage analysis. For the other, it is used to quantify novel splicing events.

*Junction support of bin coverage statistics:*. ASpli makes use of junction data as supporting evidence of alternative usage of bins. For a general differential splicing event affecting a given bin, it is always possible to define exclusion and inclusion junctions. The first class of junctions (noted as *J*_3_) pass over the bin of interest, whereas the second ones (note as *J*_1_ and/or *J*_2_) quantify and support the inclusion of start and/or end bin boundaries in the mature transcript. Panel C of Figure S8 illustrates this point for the different types of splicing events that could affect a given bin. ASpli considers for this analysis junctions that are completely included within a unique gene and have more than a minimum number of reads supporting them (by default this number is five).

PSI (percent spliced in) [45] and PIR (percent of intron retention) metrics are two well known statistics that can be used to quantify the relative weight of inclusion evidence for different kind of splicing events (see Panel C of Figure S8). For each bin, ASpli quantifies the inclusion strength in every experimental condition using the appropriate inclusion index (see Table S1). Only junctions that pass an abundance filter criterium (a minimum number of counts should be attained in all samples of at least one condition) are considered for the estimations.

**Table (S1).**
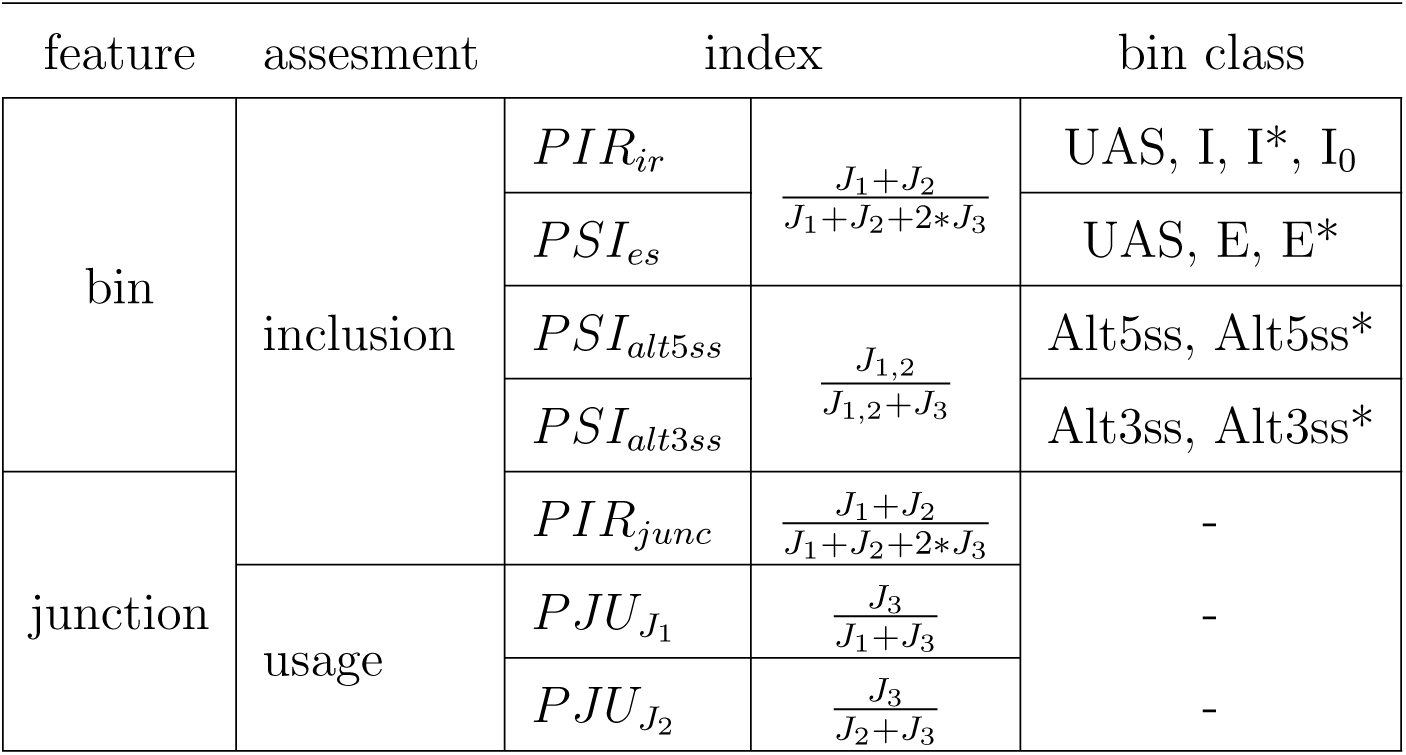
Junction usage and inclusion strength figure of merits for different bin classes and for experimentally detected junctions. The definition of *J*_1_, *J*_2_ and *J*_3_ junction counts is depicted in panels C and D of Figure S8 for annotated and experimentally detected junctions respectively.

For each bin, a PIR or a PSI metric is calculated, according to the splicing event category assigned to that bin (see last column of table S1). If no splice event was assigned, meaning that the bin is not alternative, an exon will be considered to be involved in a putative exon skipping splicing event, and an intron will be considered to be involved in a putative intron retention splicing event.

*Novel and non-canonical splicing patterns:*. ASpli relies on the direct analysis of experimentally observed splicing junctions in order to study novel (i.e. non-annotated) splicing patterns.

For every experimental junction, ASpli characterizes local splicing patterns considering two hypothetical scenarios. For one hand, assuming that every detected junction might be associated to a possible intron that could be potentially retained, a *PIR*_*junc*_ value is computed (panel D of Figure S8).

On the other hand, every junction also defines potential 5’ and 3’ splicing sites. It can be the case that one (in an alternative 5’ or 3’ scenario), or both ends (in case of exon skipping) were shared by other junctions. In this context, it is informative to characterize the relative abundance of the analyzed junction (dubbed *J*_3_) with respect to the locally *competing* ones. ASpli estimates *percentage junction-usage* indices, 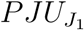 and 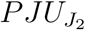, in order to evaluate and quantify this quantities (see Panel D of figure S8 and Table S1). In order to illustrate this point, we show in Panel E of figure S8 an hypothetical splicing scenario for a given junction of interest, *J*_3_. It can be appreciated that 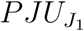 quantifies the participation of this junction in the context of a splicing pattern involving the two orange competing junctions, whereas 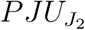 reports on the usage of *J*_3_ in connection with the green competing junction.

### 8.2. Command-line running arguments

Command lines used to invoked algorithms and further calculation details:

- STAR aligner For PRMT5 datasets

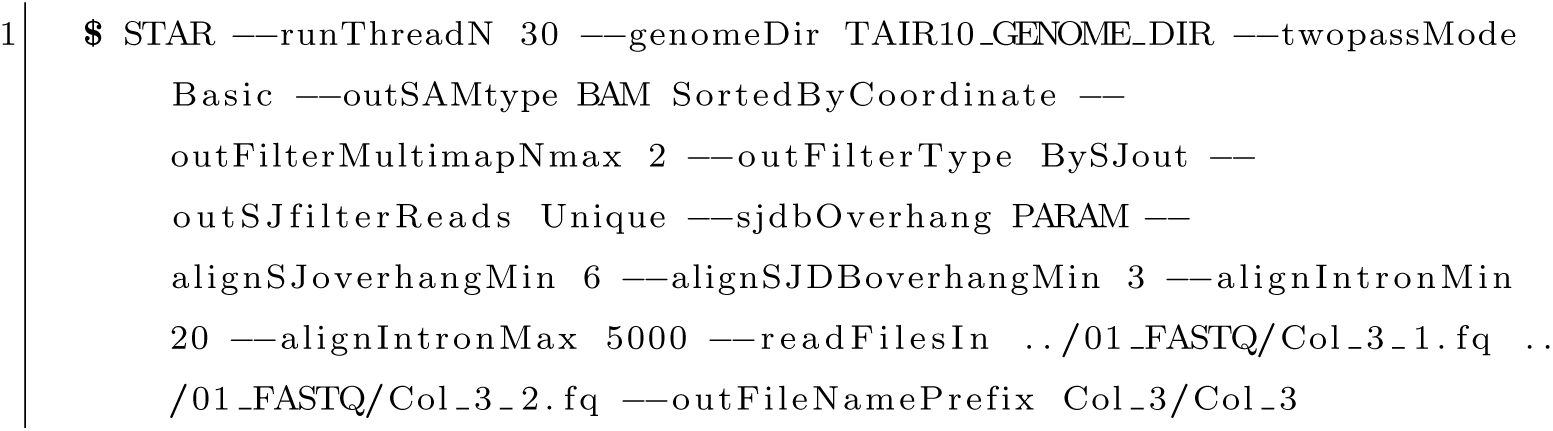 We used a sjdbOverhang parameter value equal to 99 and 149 for PRMT5 datasets A and B respectively. For the prostate dataset we aligned using default STAR parameters.

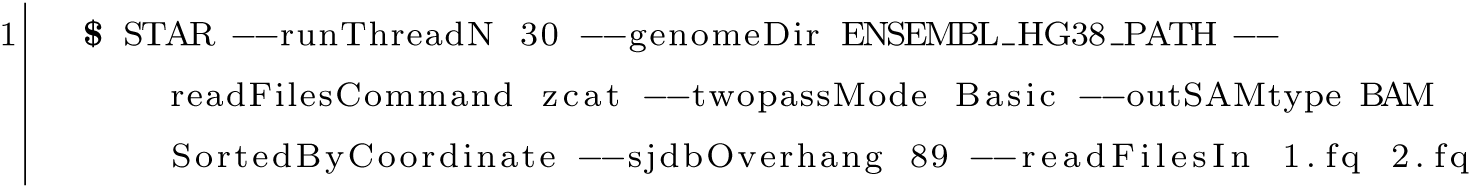 We used a sjdbOverhang parameter value equal to 99 and 149 for PRMT5 datasets A and B respectively.
- LeafCutter (synthetic dataset) BAM files were first processed using the provided *bam2junc.sh* script. The generated *juncfiles.txt* was then used to build junction clusters via the provided python script

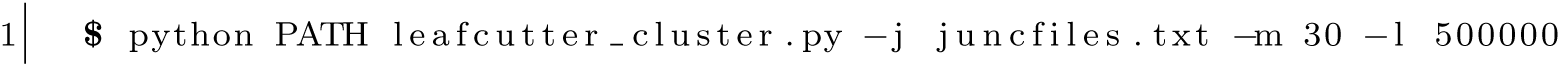 Finally, we used the provided *leafcutter ds* R-script to run the statistical analysis (min samples per intron=3).
- rMATS Command line use to analyzer PRMT5 assays:

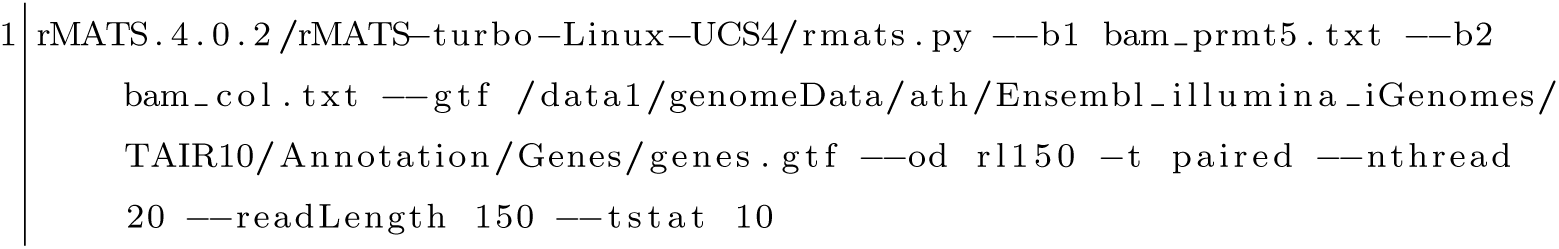
- MAJIQ

### 8.3. Splicing affected regions detected by different algorithms

Each algorithm reports splicing altered genomic features in different ways. In order to standardize the identification of regions of interest we proceeded as follows:

- LeafCutter: We first identified clusters presenting adjusted pvalues*<* 0.05 as reported in ‘leafcutter ds cluster significance.txt’ file. For each of these statistically significant clusters we considered the associated genomic-regions reported in ‘leafcutter ds effect size.txt’ file with |ΔΨ| *>* 0.1.
- MAJIQ: We considered the genomic-region covering junction clusters presenting at least one junction with *P* (|ΔΨ| *>* 0.2) *>* 0.95.
- rMATS: We considered the values reported in ‘JCEC.txt’ files. This means that we considered a model that evaluated splicing with reads that spanned splicing junctions and reads on targets bins (i.e. alternatively spliced exons). We kept junctions presenting adjusted FDR*<* 0.0 and inclusion signal larger than a 0.1 level. Genomic regions were then defined according the following rules:
- A3SS’ (A3SS.MATS.JCEC.txt file): We considered the genomic region between ‘shortEE’ and ‘longExonEnd’ coordinates for negative strand and by ‘longExonStart 0base’ and ‘shortES’ for positive strand cases.
- A5SS’ (A5SS.MATS.JCEC.txt file): We considered the genomic region between ‘shortEE’ and ‘longExonEnd’ coordinates for positive strand and by ‘longExonStart 0base’ and ‘shortES’ for negative strand cases.
- MXE (MXE.MATS.JCEC.txt file): We considered two regions per event defined by: ‘1stExonStart 0base’, ‘1stExonEnd’ and ‘2ndExonStart 0base’, ‘2ndExonEnd’.
- SE (SE.MATS.JCEC.txt file): We considered the regions between ‘exonStart 0base’ and ‘exonEnd’.
- RI (RI.MATS.JCEC.txt file): We considered the regions between ‘riExonStart 0base’ and ‘riExonEnd’.

### 8.4. Analysis of false positive calls in simulated dataset

In our simulations a 20% level of random variability was added to variant concentration profiles. A splicing activation signal (SAS) value was then estimated for each gene as the maximum absolute change in variant concentration observed between conditions. The left-most first and second boxplots in Figure S9 depict the distribution of this quantity for the 915 genes for which a splicing event was simulated, and for the remaining 7518 genes respectively. On the other hand, the four right-most boxplots show the SAS distribution for false positive calls obtained with different methods. Non explicitely splicing simulated changes were reported for 9, 4, 48 and 23 genes according to ASpli, LeafCutter, MAJIQ and rMATS algorithms respectively.

**Figure (S9).**
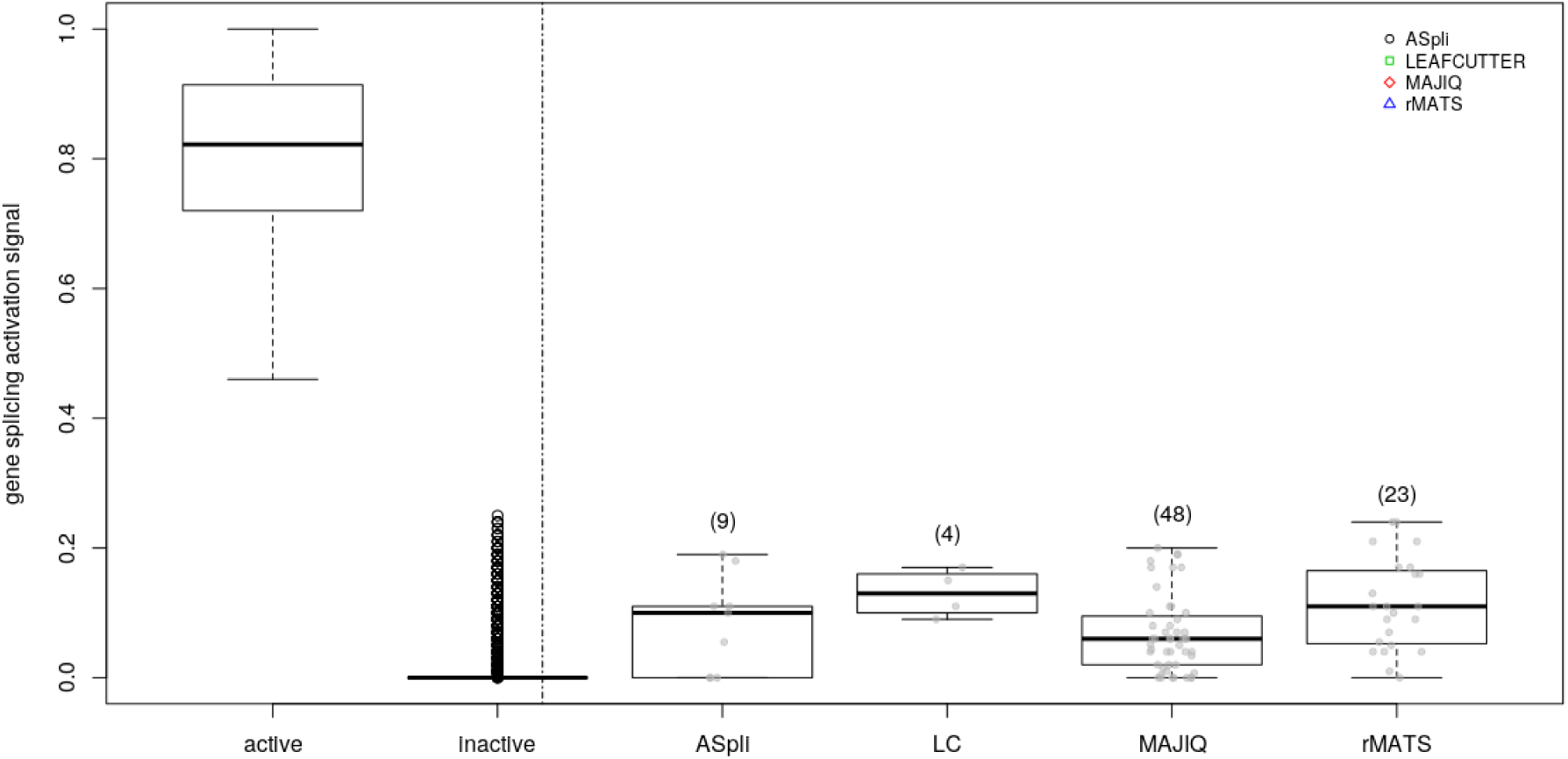
Splicing simulation.

### 8.5. Comparison of discoveries

A comprehensive comparison of discoveries appeared at first-sight problematic as each algorithm is focused on different genomic features in order to chart splicing landscapes.

For instance, rMATS analyzes genomic regions flanked by upstream and downstream exons to examine canonical splicing events. MAJIQ and LeafCutter, on the other hand, exclusively rely on clusters of split reads that share start or ending junction-ends. Finally ASpli considers both, junction clusters and bin features, i.e. genomic regions defined from disjoint ranges of annotated junctions.

In this context, a first coarse grained comparison could be established at gene-level, comparing the identity of genes housing splicing-altered patterns according to the different analyzed methods. Panel (A) of Figure S10 displays a color-coded overlap matrix of affected genes in experiments *A* and *B* according to the four examined methodologies. Each cell reports the intersection size and, in brackets, the corresponding overlap coefficient. At gene level, rMATS achieved the largest agreement factor (83% of genes identified in experiment *B*, were also reported in experiment *A*). However, it also produced the lowest number of discoveries (119). ASpli, on the other hand, presented a comparable level of agreement (71%), highlighting a significatively larger number of concordant genes (2109). Typically, more than 50% of genes identified by any methodology was also reported by ASpli (first and second rows of Figure S10). Moreover, the number of concordant discoveries between experiments considering a given methodology was comparable to the agreement level achieved between each experiment-metodology combination and the correpsonding ASpli result. Noticeably, more than 90% of MAJIQ’s genes were also spotted by ASpli.

A more in-depth comparison could be established analyzing the overlap of identified genomic regions. In panels (b) and (c) of Figure S10 we informed the extent of the overlaps between genomic regions found to be affected by differential splicing patterns according to each algorithm (see Material and Methods 3.5) to map events reported by each method to a common set of genomic coordinates). While any kind of overlap was registered for panel (b), only complete inclusion of genomic regions identified by one method inside the ones identified by a second one was considered for panel (c). Statistically significant overlaps were marked with asterisks. Note that overlap coefficients (in brackets) exceeding unity were detected in between-experiments comparisons for LeafCutter and rMATS as a result of the presence of one-to-many region mappings.

**Figure (S10).**
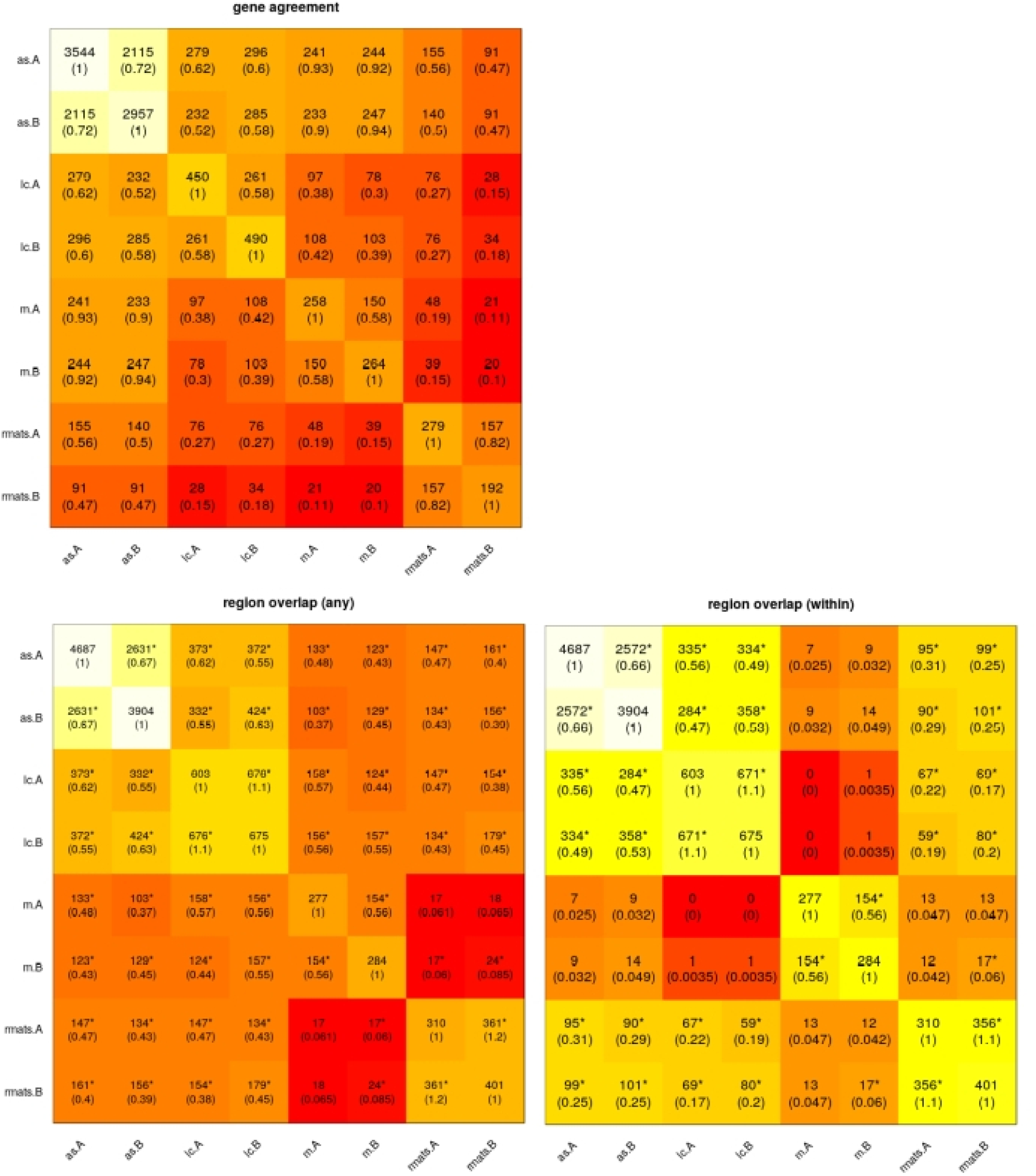
ASpli main functions.

For the loose overlap criterium we found statistically significant concordance between discoveries for almost every cell (Fig S10-b). Only specific comparisons involving MAJIQ and rMATs failed the statistical significance test. At the same time, overlap coefficient values were similar to the ones estimated at the gene-level analysis. Noticeably, we recognised a sensible reduction in this quantity for the MAJIQ vs ASpli comparison. This finding highlighted that gene-level agreement should in general be considered with caution. A more detailed examination at the sub-genic level might be necessary to assess for discovery consistencies between algorithms. Results for the most stringent overlap criterion are shown in Figure S10(c). As expected, a major decrease on overlap coefficient values was observed. However, statistically significant agreement between results was still found as a general rule. Only comparisons involving MAJIQ’s discoveries failed the statistical assessments.

### 8.6. PRMT5 PCR events

We characterized the agreement between the 23 splicing events that ASpli uncovered for the consolidated AB case, and the 44 Sanchez qRT-PCR validated events in Table S2. For each assayed event we included the kind of the original event and the reported qRT-PCR splicing signal value in the second and third columns respectively (Sanchez and collaborators calculated the fraction of the shortest isoform in PRMT5 mutants and wildtype plants detected by qRT-PCR, and used the relativized difference between them as a quantitative proxy of splicing changes (Table 4 of [25])). In the fourth column we informed whether the PCR-interrogated genomic region overlapped with the one signaled by ASpli. Finally, the type of splicing event detected by ASpli was included in the last column of the table.

### 8.7. Prostate cancer dataset:Transcriptomic variability

In order to visualize the transcriptomic variability across patients at gene expression levels we considered the 30% most variable genes across the 28 expression profiles that presented more than 10 counts per million reads in at least 3 samples. With this informative set of 1386 genes we built a multidimensional scaling plot of distances between gene expression profiles estimated with the edgeR package [24]. Results are shown in Fig S11. In this kind of plot, samples lay on a two-dimensional scatterplot so that distances on the plot approximate the typical log2 fold changes between the samples (function plotMDS of edgeR [24]).

Emtpy and filled symbol correspond to tumor and normal tissue samples respectively. Pair of points of a given patient are equally colored and joined by a dashed edge.

It can be seen that tumor and normal samples were well separated across the leading reduced dimension. The second largest projected dimension, however, let us appreciate internal structure and some variability between patients. There was a group of 5 patients (top left empty points) that displayed a rather homogeneous pattern of changes between tumor affected and normal tissues. On the contrary, the 9 bottom-left tumor samples seemed to segregate into a different cluster of transcriptomes. Moreover, the corresponding patients presented different kinds of alterations between tumor and control samples.

**Figure (S11).**
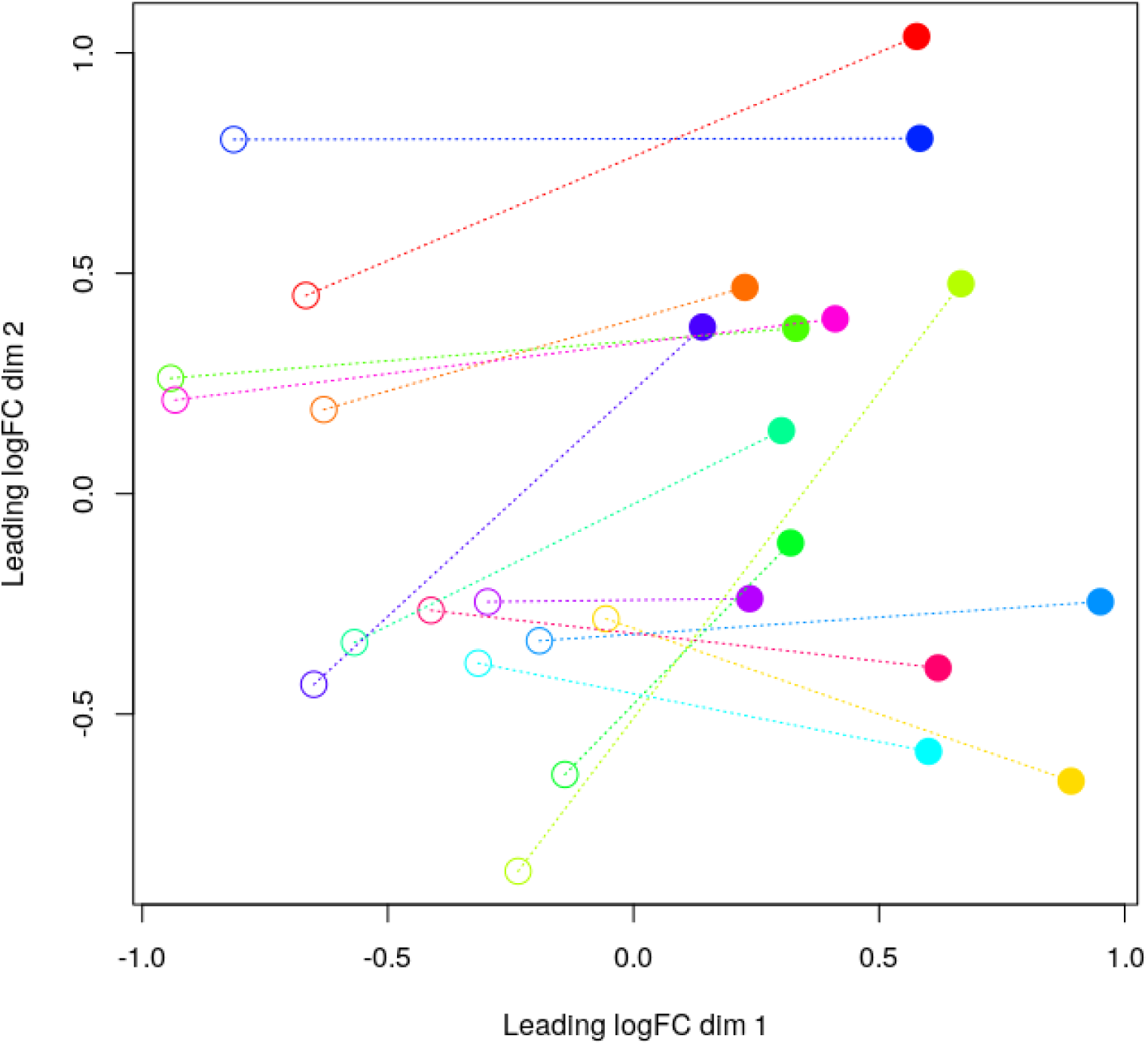
A.

**Table (S2).**

|  | Gene ID | Event | qRT-PCR signal | Region overlap | Detected event |
| --- | --- | --- | --- | --- | --- |
| 1 | AT1G53650 | 5'ss | 18.93 | yes | IR (next to 5) |
| 2 | AT1G54360 | 5'ss | 39.21 | yes | Alt5ss |
| 3 | AT1G76510 | 5'ss | -28.00 | yes | Alt5ss |
| 4 | AT2G04790 | 5'ss | 11.14 | yes | IR |
| 5 | AT2G15530 | 5'ss | 21.12 | yes | Alt 5'/3' |
| 6 | AT2G33480 | 5'ss | -27.16 | yes | IR |
| 7 | AT2G38880 | 5'ss | -10.44 | no | IR |
| 8 | AT2G46790 | 5'ss | 35.20 | yes | Alt5ss (plus additional IR) |
| 9 | AT3G01150 | ES | -13.70 | no | IR |
| 10 | AT3G12250 | 5'ss | -16.29 | no | ES* |
| 11 | AT3G16800 | IR/3'ss | -31.59 | yes | IR,Novel Alt 5'/3' |
| 12 | AT3G19840 | 5'ss | -26.20 | no | IR |
| 13 | AT3G20270 | ES | 8.51 | no | IR (next to ES) |
| 14 | AT3G23280 | ES | 18.21 | yes | ES |
| 15 | AT3G25840 | ES | 6.51 | yes | ES |
| 16 | AT4G02430 | 3'ss | 27.45 | no | IR |
| 17 | AT4G24740 | ES | 16.07 | no | IR (next to ES) |
| 18 | AT4G31720 | 3'ss | 12.37 | no | IR |
| 19 | AT4G32730 | 5'ss | 30.93 | yes | Novel Alt 5'/3' |
| 20 | AT4G38510 | 5'ss | 15.53 | yes | Alt5ss*,CSP |
| 21 | AT5G05550 | ES | 32.72 | yes | ES (plus additional IR) |
| 22 | AT5G25610 | IR | -71.69 | yes | IR |
| 23 | AT5G57630 | 5'ss | 31.07 | yes | Novel Alt 5'/3' (plus adjacent IR) |

